# Mechanical strain and strain rate define thresholds of single-cell and ensemble calcium disruption in traumatic brain injury

**DOI:** 10.64898/2026.11.20.768182

**Authors:** Jamie A. Sergay, Natalie Schick, Emily Blick, Mauricio Araiza Canizales, Jing Zhang, Christian Franck, Aviad Hai

## Abstract

Mechanically induced brain injury triggers rapid cellular responses that shape long-term neurological outcomes, yet the early evolution of functional network disruption remains poorly defined. In particular, how neuronal activity changes within seconds of deformation and whether these responses determine subsequent circuit organization remain unknown. Here, we combined controlled mechanical stretch with high-speed calcium imaging to measure neuronal calcium dynamics within seconds of injury and track the same cells over 24 hours. We identify strain- and strain-rate-dependent disruption modes, including rapid reduction of calcium transient activity under moderate loading and collapse of network bursting beyond distinct mechanical thresholds. These disruptions persist despite preserved cell viability, revealing an additional functional injury pathway. These findings show that functional impairment emerges within seconds of mechanical loading and that the severity of deformation dictates the mode of network disruption, establishing mechanical thresholds as key determinants of secondary injury progression.

**Significance:** Mild traumatic brain injury often impairs brain function without overt tissue damage, but how the mechanical insult itself disrupts neuronal communication in the first seconds after impact has been difficult to measure. By combining precisely controlled stretch of cultured neural networks with high-speed calcium imaging, we resolved neuronal activity within seconds of deformation and tracked the same cells for 24 hours. Strain magnitude and strain rate set distinct thresholds for network disruption, ranging from graded suppression of activity to collapse of synchronized bursting and altered calcium homeostasis, all without loss of cell viability. These mechanical thresholds quantitatively link tissue deformation to the earliest functional injury and can inform injury prediction and preclinical models.

## Introduction

Traumatic brain injury (TBI) spans a complex spectrum of severity, involving biochemical and electrophys-iological cascades that can lead to the disruption of neuronal network dynamics, acute cognitive deficits, and potential chronic neurodegeneration (1–6). Mild TBI (mTBI), the most common form of TBI (7), produces unpredictable alterations that include irregular excitation and inhibition (8–11), dysregulated networks (8, 12–15), and reduced local and global brain connectivity (15–17). These changes compromise information integration, which is essential for cognition, and may contribute to disease pathology while also informing therapeutic strategies (14, 18, 19). However, uncovering the direct relationship between the primary mechanical insult in mTBI and the resulting functional disruptions has been challenging. This is mainly due to the limited understanding of how specific mechanical loading conditions give rise to clinical etiology. It also reflects the current constraints of experimental systems capable of applying precise, reproducible perturbations while simultaneously monitoring large-scale neuronal network activity relevant to mTBI (20–22). Establishing links between specific mechanical loading conditions and modes of network alteration can help explain the signaling disruptions that underlie mTBI pathophysiology and can potentially resolve the functional basis of the earliest clinically relevant impairments such as brief loss of consciousness (23).

Tissue strain and strain rate have emerged as key biomechanical determinants of injury pathology. Head impacts generate propagating strain fields that deform neural cells, triggering secondary injury cascades and cognitive decline (12, 24–29). Injury magnitude modulates electrophysiological signaling patterns and connectivity within mechanosensitive networks (21, 30, 31). Variations in strain rate produce distinct cellular pathomorphological outcomes (26, 29, 32) and time-dependent synaptic deficits (33) that are coupled to specific molecular pathways regulating neuronal excitability (34). Mechanosensitive channels may help explain how deformation is converted into early functional disruption (30). Mechanically gated NMDA receptors and TRPV4 provide plausible routes for stretch-associated calcium influx (35–37). Piezo-family channels may also contribute to mechanically evoked calcium signaling in neural tissue (38), whereas mechanosensitive potassium channels such as TREK-1 and TRAAK can suppress neuronal excitability during membrane stretch (39). Together, these pathways suggest that the same loading event could differentially alter spike-coupled calcium transients and basal calcium state after mTBI. While strain and strain rate reliably predict cellular damage, their precise influence on network-scale electrophysiology is not fully understood (8, 40, 41). Cellular-level studies have more commonly focused on understanding the initiation of cell death, often overlooking minute electrophysiological perturbations that are more relevant to mTBI. However, subtle disruptions can impair executive function at strain thresholds below those inducing cell death (42). Network-level readouts that can be characterized in parallel with cellular injury and cell death are therefore essential for capturing the temporal evolution of post-injury deficits.

Here, we combine high-resolution calcium imaging across multiple time scales with controlled mechanical loading to establish a parametric link between mechanical deformation and calcium transients, homeostasis, and cell viability at single-cell resolution across large neuronal populations. To examine deformation relevant to mTBI, we systematically vary applied strain (0.1–0.5) and strain rate (1–50 s*^−^*^1^) using a precision-controlled tension platform, encompassing validated thresholds for electrophysiological and structural sequelae (24, 27, 43, 44). Analyses reveal nonlinear, strain- and rate-dependent differences in calcium transient kinetics, network participation, and ionic homeostasis. We also dissociate functional disruption from cell death at these loading regimes. This study captures major signaling alterations at the hyperacute stage (seconds to minutes) post-injury that are relevant to the molecular machinery. This approach overcomes limitations of benchmark experimental paradigms (31, 43, 45), which typically observe changes only minutes to hours after insult due to challenges in coupling mechanical deformation with rapid functional measurements. By quantifying calcium dynamics within seconds of injury and tracking neuronal populations over 24 hours, we establish a mechanistic framework that links mechanical deformation to the early disruption of neuronal networks during the hyperacute phase of brain injury and to its subsequent evolution. This system identifies mechanistically grounded targets for injury prediction, monitoring, and neuroprotective intervention.

## Materials and Methods

### Primary cortical neural cell isolation and culture

Cortical neural cell cultures were acquired from female postnatal Sprague-Dawley rats (p0-p1). All animal procedures were conducted in accordance with the guidelines approved by the University of Wisconsin-Madison Institutional Animal Care and Use Committee (protocol M006019-R02-A02). All cortical tissue was isolated from female pups; no male animals were used in this study. Culture substrates were first prepared in the shape of dog-bones using the protocol presented in Summey et al. (46). Briefly, a mixture of polydimethylsiloxane (PDMS; SYLGARD 184, 10:1 base-to-curing agent; Dowsil) was molded into dog-bone shapes using SU8 molds on silicon wafers and then autoclaved to sterilize. The sterile PDMS dog-bones were treated with a mixture of laminin (4 *µ*g/mL; Thermo Fisher Scientific) and poly-D-lysine (0.1 mg/mL; Thermo Fisher Scientific) for 60 minutes at 37 *^◦^*C, then washed three times with PBS (1X; Thermo Fisher Scientific).

Cortical tissue was isolated, and cells were dissociated following the protocol by Scimone et al. (47). Briefly, the cortices are isolated and processed in papain solution (Worthington Biochemical). Cells were cultured in a complete cortical media (CCM) consisting of Neurobasal-A medium, 2% (v/v) B-27 Plus supplement, 1.01% (v/v) penicillin-streptomycin, 0.87% (v/v) DMEM, 0.25% (v/v) GlutaMAX, 0.1% (v/v) fetal bovine serum, and 0.02% (v/v) L-glutamine (all components from Thermo Fisher Scientific). Cells were seeded at 28,000 cells/sample and maintained in a humidity-controlled 37 *^◦^*C incubator with 5% CO_2_. Cultures were grown for 20 days in vitro (DIV), and media was replaced every 2-3 days, alternating between partial and full media exchanges.

To characterize the cellular phenotypes in the neural cultures, samples were immunolabeled for *β* -III Tubulin and glial fibrillary acidic protein (GFAP), markers of neurons and astrocytes, respectively (Fig. 1B). At DIV 20, cultures were fixed with 4% paraformaldehyde (Thermo Fisher Scientific) for eight minutes and permeabilized with 0.5% (v/v) Triton X-100 (Sigma Aldrich). To reduce nonspecific antibody binding, samples were blocked with 10% BSA for 30 minutes and washed twice in 0.5% BSA (Sigma Aldrich). Cultures were incubated for 1 hour at room temperature in a primary antibody solution containing mouse anti-*β* -III Tubulin (1:500, Invitrogen Cat#PA5-85639, RRID:AB_2792779) and chicken anti-GFAP (1:500, Abcam Cat#ab4674, RRID:AB_304558) diluted in 0.5% BSA. Following four washes in 0.5% BSA, cultures were treated with a secondary antibody solution for one hour at room temperature that contained a dilution of goat anti-mouse AlexaFluor 555 (1:500, Invitrogen Cat#A21422, RRID:AB_141822) and goat anti-chicken AlexaFluor 488 (1:500, Invitrogen Cat#A11039, RRID:AB_2534096) in 0.5% BSA. Samples were subsequently washed twice with 0.5% BSA and 1X PBS, then stored in PBS at 4 *^◦^*C until imaging. All images were collected using a Nikon Ti-E confocal microscope (RRID:SCR_021242) at 20X magnification and processed in Fiji (RRID:SCR_002285).

**Figure 1.**
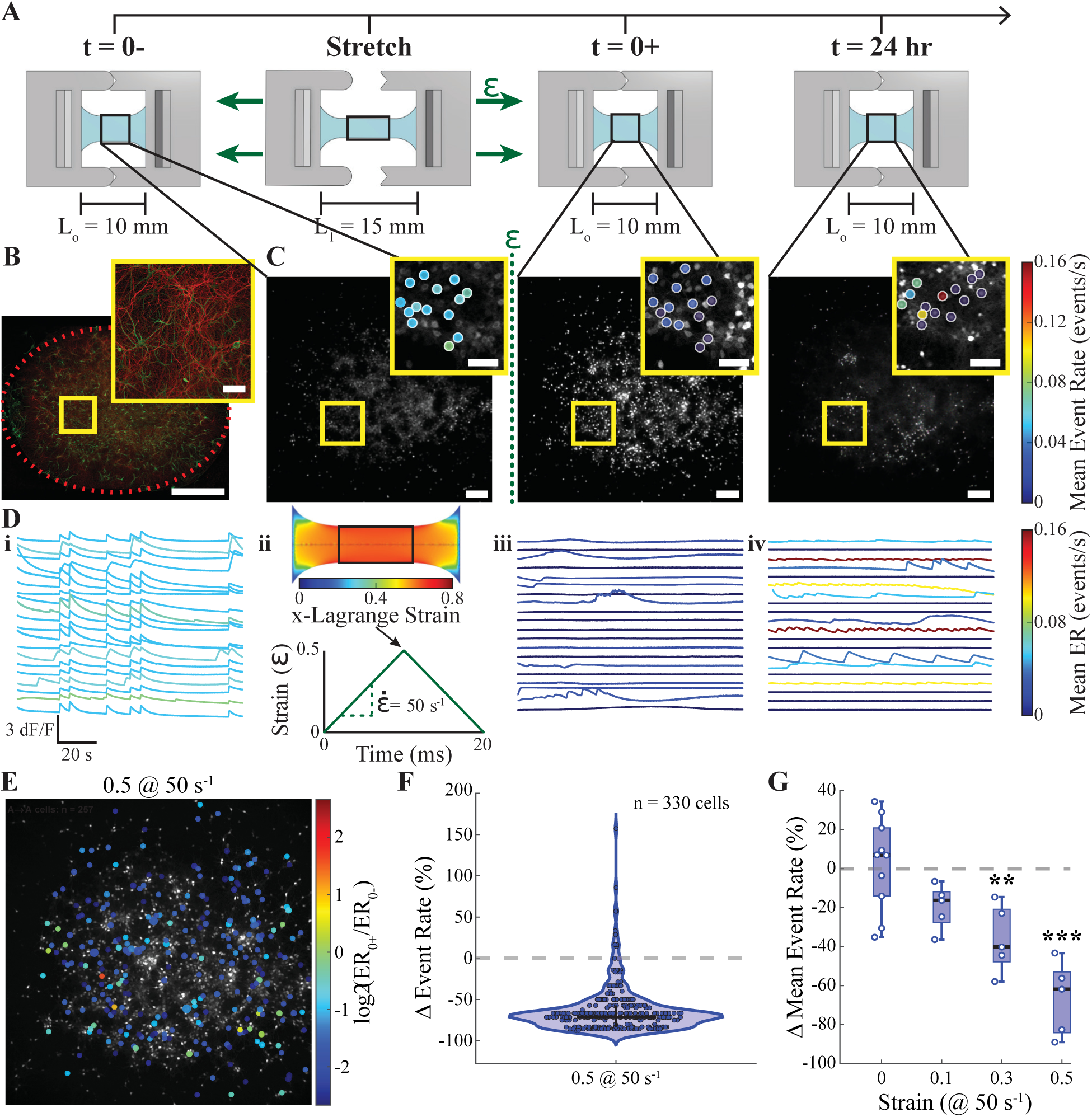
Mechanical stretch causes immediate hypoactivity in neuronal networks. (**A**) Recording timeline and setup: PDMS “dog-bone” substrate in a uniaxial stretch platform; seeding region highlighted. (**B**) Phenotypic makeup of 20 days in vitro (DIV) neuron-astrocyte co-cultures (*β* III-tubulin, red; GFAP, green). Scale bar, 1 mm (inset, 100 *µ*m). (**C**) hSyn1-GCaMP6s images from the same FOV immediately before stretch (0-), immediately after (0+), and 24 hours post-stretch (24h); color encodes per-cell event rate (ER). Scale bar, 200 *µ*m (inset, 100 *µ*m). (**D**) Temporal experimental profile: (i) representative calcium traces at 0-, (ii) loading waveform (*ε* = 0.5 at 50 s*^−^*^1^) with dog-bone schematic showing the Lagrange strain field at peak strain with the seeded region highlighted, and traces at (iii) 0+ and (iv) 24h; color encodes ER. (**E**) Spatial map of a representative 0.5 strain @ 50 s*^−^*^1^ field colored by *log*_2_(*rate*_0+_*/rate*_0_*_−_*) (jet color map; A*→*A cells only, active at both recordings). (**F**) Per-cell distribution of the immediate change in ER (0+ vs 0-) from the representative cell population in (E) showing a downward-shift consistent with broad per-cell slowing (violin + beeswarm; n = 300 cells; A*→*A cells). (**G**) Immediate network-level ER percent difference (0+ vs 0-) shows graded strain dependence at 50 s*^−^*^1^ (box plots; *N* = 4 *−* 10 cultures per condition; \*\**p <* 0.01, \*\*\**p <* 0.001).

Culture composition was estimated by comparing the number of GCaMP6s-positive somas (neurons; FIJI Analyze Particles) to the total number of Hoechst-positive nuclei (all cells; custom MATLAB counter) in matched fields of view immediately before stretch. Across analyzed samples, neurons accounted for 55.3 *±* 20.3% of total cells (mean *±* SD, n=42; three samples were excluded due to modality discrepancies in edge-intensity correction during segmentation). This neuron fraction is modestly enriched relative to the *∼*40% neuron fraction reported for adult rat cerebral cortex (48) and consistent with the developmental stage of the cells and the neuron–astrocyte co-culture composition supported by the maintenance media.

### Uniaxial tension model

A custom-built uniaxial tension device was used to apply programmed, single-pulse stretches to the PDMS dog-bone samples (46). The stretch sequence is illustrated in Fig. 1D. The applied strain was previously validated in Summey et al. on the same PDMS substrate and geometry used here (46). The experimental cell cultures were stretched at strain rates of 1 strain/sec (s*^−^*^1^) or 50 s*^−^*^1^ and strains of 0.1, 0.3, or 0.5. The tension device was mounted atop a Nikon Ti-E microscope stage to enable immediate imaging before and after the stretch.

The strain field in the cell culture region was visualized using finite element simulations. The half-plane symmetry dog-bone geometry was recreated in SolidWorks 2024 and imported as an STL file into FEBio Studio 2.10.0 (RRID:SCR_023056) for modeling. Tet4 and a size of 0.07 were applied to all elements in the model. For an applied strain of 0.5, it was imperative to use a material model that captures PDMS behavior under such extreme conditions. An Ogden 3 material model fits the experimental data best for 250*µ*m thick PDMS films stretched to 0.5 (49). The bulk modulus was kept at the default value of 50 MPa to simulate incompressibility. The final geometry and mesh, as well as the displacement boundary conditions are visualized in Fig. S1. Briefly, four boundary conditions were applied to the surfaces of the geometry to mimic the experimental conditions. (I) A symmetry boundary condition was applied along the half-symmetry plane surface to prevent out-of-plane motion in the y- and z-directions. (II) A displacement boundary condition of 2.5 mm in the x-direction was prescribed to the right most face to model the stretch in the positive x-direction. (III) A similar boundary condition but with a prescribed displacement of -2.5 mm in the x-direction was applied to left most face to stretch the geometry in the negative x-direction. (IV) A final boundary condition consisting of fixing the displacements in the y- and z-direction was applied to the latter two faces to mimic the fixed ends of the dog-bone.

### Fluorescent imaging

All images were captured using widefield imaging with a Nikon Ti-E microscope, 10X objective (0.45 NA; Nikon), and a high-speed CMOS camera (Teledyne Kinetix). All time-lapses had a 1600×1600 pixel field of view (FOV). Spontaneous calcium activity was acquired at a nominal rate of 25 Hz with 12-bit, 2×2 binning, 8% LED intensity, and 3-5 ms exposure times using a standard FITC filter cube. Two-minute time-lapses were recorded immediately before stretch (0-), 2.5 seconds (95% CI [1.7, 3.3] s) post-stretch (0+), and 24 hours post-stretch (24h). At 24h, the samples’ FOV was matched to the calcium image at 0- within a few pixels. Images for cell viability were captured from the same samples at 16-bit, 3200×3200-pixel FOV, 8% LED intensity, and 65 ms exposure time using standard DAPI and TRITC filter cubes. The microscope stage was enclosed in a black-out temperature-control box set to 37 *^◦^*C throughout all experiments. Imaging parameters were set to prevent fluorescent oversaturation, even after stretching.

### Neuronal calcium transient assessment

Neural network activity was quantified by optically capturing the calcium dynamics during spontaneous network activity. Cultures were transfected with neuron-specific GCaMP6s at 7 DIV (AAV1-Syn-GCaMP6s-WPRE-SV40; from Douglas Kim & GENIE Project; Addgene plasmid #100843, RRID:Addgene_100843). Cells were transfected at a concentration of 5 *×* 10^10^ GC/mL for 48 hours, with warm CCM added 24 hours into the process.

Calcium fluorescent signals were pre-processed with NeuroCa (50). From the GCaMP timelapses, somas were detected, fluorescent intensity traces extracted, corrected for photobleaching, and normalized. All other signal processing and quantification were performed according to the protocol in NeuroSpikeX (51). After defining the noise floor, calcium transient peaks (events) were detected, and their decays were analyzed. The event rate was calculated per cell and per sample for all active cells (*≥* 1 event). All plots showing percent change in event rate (ER) were calculated as ΔER = 100 *×* (ER_0+_ *−* ER_0_*_−_*)*/*ER_0_*_−_* or ΔER = 100 *×* (ER_24*h*_ *−* ER_0_*_−_*)*/*ER_0_*_−_*. Synchronous network events (bursts) were automatically calculated as a continuous period during which at least 20% of active cells had an event. To calculate the decay rate, each transient was shifted so that the peak started at *t* = 0 and the decay ended at *dF/F* = 0. Then, the exponential decay, *dF/F* = *Ae^−kt^*, was fitted, and the rate constant, *k*, was averaged per-cell and per-sample.

The relative basal fluorescence intensity shift was also calculated using NeuroSpikeX. Frames from the 0-and 0+ time-lapses were compared immediately before and after the stretch, where most cells were at their resting state. To classify meaningful changes, a threshold was set by calculating *log*_2_(Intensity_0+_*/*Intensity_0_*_−_*) for all tracked neurons in the sham samples. The 95th percentile of values was classified as unchanged; values above the threshold were considered an elevated fluorescence, and values below the threshold were considered a reduced fluorescence. This classification was applied to all stretch conditions.

Prior to analysis, all samples underwent four quality control checks. First, neuronal cell density was estimated from the GCaMP6s fluorescence signal in FIJI; samples with fewer than 150 neurons/mm^2^ were excluded to ensure cultures were sufficiently dense for network-level analysis. The final analyzed samples had densities 468 *±* 222 neurons/mm^2^ (mean *±* SD). Second, the network-level calcium event rate at 0- was required to be *≥*0.01 events/sec; samples below this threshold were excluded as they were considered insufficiently active at baseline to assess injury-induced changes. Third, samples with insufficient baseline calcium activity for reliable event detection and analysis were excluded. Fourth, mean background-subtracted resting fluorescence at 0- was compared across all samples to confirm no preparations had anomalous signal levels that could distort dF/F-based measurements. A total of 13 samples were excluded prior to analysis: 7 due to low neuronal density, 3 due to insufficient baseline calcium activity, 1 due to low network event rate, and 2 due to experimental handling errors during data collection.

### Cell viability assay

Cell viability after 24 hours post-stretch was determined by staining cultures with Hoechst 34580 (Thermo Fisher Scientific) and Ethidium Homodimer 1 (Thermo Fisher Scientific). All staining and imaging media were warmed to 37 *^◦^*C prior to use. Samples were washed once with Hanks’ Balanced Salt Solution (HBSS; Thermo Fisher Scientific). The Hoechst solution (10 *µ*M in HBSS) was added to the samples for 10 minutes before the 0- image, then washed out and replaced with the imaging media. Imaging media was buffered for ambient CO_2_ levels by adding 15 mM HEPES (Thermo Fisher Scientific) to CCM. Just prior to the 24h imaging, the ethidium solution (2 *µ*M in HBSS) was added for 10 minutes and washed out. Control samples that were never placed in the tension device nor imaged at 0-/0+ were stained with both Hoechst and ethidium for 10 minutes at the 24-hour time point.

Cell death was calculated for each sample in MATLAB (RRID:SCR_001622) by first transforming the Hoechst capture at 24 hours to match the Hoechst capture at 0-. The same transformation was applied to the ethidium image. The Hoechst signals were tracked between the two time points. To account for any small cell drifts, any percentage of signal overlap, and a signal within 9 pixels (*≈* 1*/*2 median nuclear diameter) was counted as a surviving cell. All other lost Hoechst signals at 24h were classified as dead. Cell death was further refined by counting any cells co-localizing ethidium with Hoechst at 24h as dead, as this indicates a compromised membrane.

### Experimental design and statistical analysis

Cultures were randomly assigned to strain and strain-rate conditions, and the order in which conditions were run was varied across experiment days so that day-to-day variability was not confounded with a specific condition. At least six independent cultures were planned per condition based on convention rather than an a priori power calculation. Thirteen cultures were excluded prior to any group comparison according to the quality-control criteria described above, yielding the *N* = 4–10 independent cultures per condition reported in the Results and figure legends; no cultures were excluded after outcome data were inspected. Investigators were not blinded to condition during stretch delivery and image acquisition, since the experimenter directly set the applied strain and strain rate. All downstream quantification (event detection, burst classification, basal-fluorescence classification, and viability classification) used automated pipelines applying identical, condition-independent parameters to every sample. Sample sizes reflect the number of independent cultures (biological replicates), not individual cells. This study was not preregistered.

Statistical analysis was performed in R 4.5.1 (RStudio; RRID:SCR_001905) using the glmmTMB and emmeans packages. Population-level metrics that were non-negative and could include true zeros (calcium event rate, event intensity, network burst rate, and number of active cells) were analyzed using generalized linear mixed models (GLMMs) with a Tweedie distribution and log link. Transient decay rate constants, which are strictly positive, were analyzed with a Gamma GLMM and log link. Proportion outcomes with known denominators (neurons with elevated basal fluorescence at 0+ relative to matched neurons, and dead cells relative to the baseline nuclear count) were analyzed with beta-binomial models and a logit link, and are reported as percentage-point differences on the response scale. Stretch/sham condition and time point were included as fixed effects, with a random intercept for Sample_ID where samples contributed more than one observation.

Immediate (0+ versus 0-) contrasts were obtained from a single model fitted across all conditions for each outcome. Contrasts between 24 hours and the earlier time points were evaluated within each condition using separate models, because Tweedie dispersion differed substantially across loading conditions and a common dispersion parameter was not appropriate; these within-condition contrasts were Šidák-adjusted across the two planned comparisons. To test whether 24-hour changes exceeded those in sham cultures, the change from baseline in each stretch condition was additionally compared with the change in sham in a model fitted across all conditions with condition-specific Tweedie dispersion (Dunnett-adjusted).

Planned contrasts and post hoc comparisons were performed using estimated marginal means. Pairwise comparisons among experimental groups were Tukey-adjusted, comparisons between experimental conditions and sham were Dunnett-adjusted, and comparisons of the two strain rates within a matched strain magnitude were planned and unadjusted. Because these are asymptotic model fits, tests are reported as Wald *z* statistics with exact *p* values and 95% confidence intervals; degrees of freedom are not defined for these models, and sample sizes refer to independent cultures. In the 0.5 strain at 50 s*^−^*^1^ condition, network bursts were absent in every culture at 0+, producing complete separation. A Wald test is not interpretable under complete separation, so this outcome was instead evaluated with a two-sided Fisher’s exact test comparing the proportion of cultures without bursts at 0+ in this condition against sham. Functional connectivity metrics were analyzed separately using Gaussian linear mixed models and are reported with Satterthwaite degrees of freedom; full details are given in the Supporting Material. Statistical significance was defined as *p <* 0.05. The robustness of each significant finding to individual sample contributions was assessed via leave-one-out sensitivity analysis, in which each model was refit with one culture omitted at a time and the proportion of refits retaining *p <* 0.05 was recorded (see Supporting Material and Table S1).

## Results

### Controlled mechanical stretch immediately suppresses network activity in a strain-dependent manner

In order to test the functional effects of various levels of applied stretch, traumatic loading was applied using uniaxial tension, and calcium activity was recorded immediately before stretch (0-), immediately after stretch (0+), and 24 hours post-stretch (24h) (Fig. 1A, C-D). Calcium activity was measured in neuron–astrocyte co-cultures on a PDMS substrate (Fig. 1B). Neuron-specific GCaMP6s was captured in two-minute recordings from the same field of view at all time points (Fig. 1C). Mimicking tissue deformations associated with mTBI, single-pulse deformations (Fig. 1D ii) were applied at nominal strains (0.1, 0.3, 0.5) and strain rates (1s*^−^*^1^, 50 s*^−^*^1^). Finite element simulation of the stretch shows a near-uniform strain field in the recorded cell region (Fig. 1D ii; see Methods for details), consistent with prior experimental measurements (46). To assess potential effects on the cellular network arising from the tension device and experimental protocol, activity in control cultures was recorded and compared with that in sham samples. This control analysis confirms that all results presented were not due to device contact or imaging (Fig. S2; see Supporting Material).

Calcium event rate (ER), defined as the rate of calcium transient peaks per neuron, was used to quantify neuronal activity. In a representative field stretched to 0.5 strain at 50 s*^−^*^1^, spatial mapping of log_2_(ER_0+_*/*ER_0_*_−_*) revealed widespread reductions in ER across individual cells throughout the field of view (Fig. 1E). Consistent with this spatial pattern, the distribution of per-cell changes from 0- to 0+ showed a marked leftward shift, with a mean percent change of -71.4% across matched active cells (A*→*A; *n* = 300 cells) (Fig. 1F).

To determine whether this suppression generalized across the full range of strain value, group-level per-sample means were compared (Fig. 1G). Immediate hypoactivity in the first two-minutes after stretch scaled with strain at 50 s*^−^*^1^. Using a log-link Tweedie generalized linear mixed model (GLMM), the estimated mean percent change (ΔER = 100 *∗* (ER_0+_ *−* ER_0_*_−_*)*/*ER_0_*_−_*) was significantly reduced at 0.3 strain (mean ΔER = *−*38.05%, 95% CI [-53.63, -17.24]%; *N* = 5; *z* = *−*3.24, *p* = 1.2 *×* 10*^−^*^3^) and 0.5 strain (mean ΔER = *−*72.07%, 95% CI [-79.45, -62.02]%; *N* = 5; *z* = *−*8.14, *p* = 4.0 *×* 10*^−^*^16^). Thus, at rapid loading (50 s*^−^*^1^), strain magnitude strongly modulates the immediate network-level reduction and creates a graded decrease in event rate that scales with strain.

### Strain rate sets the threshold for neuronal network dysfunction

Because mechanical injury depends on both strain magnitude and loading rate, we next asked whether the strain-dependent calcium response was also sensitive to strain rate. Compared to rapid loading (50 s*^−^*^1^), slow loading (1 s*^−^*^1^) produced only modest suppression of calcium activity immediately after stretch (1 s*^−^*^1^: *ε* = 0.1, *N* = 5, *z* = *−*1.24, *p* = 0.215; *ε* = 0.3, *N* = 4, *z* = 0.59, *p* = 0.554; *ε* = 0.5, *N* = 6, *z* = *−*2.08, *p* = 0.038; Tweedie GLMM). Figure 2A shows representative rasters and spectrograms of calcium event peaks at 0- and 0+ for a *ε* = 0 (sham; no stretch initiated), *ε* = 0.5 at 1 s*^−^*^1^, and *ε* = 0.5 at 50 s*^−^*^1^ culture. At 1 s*^−^*^1^, activity is only mildly reduced, whereas at 50 s*^−^*^1^ activity is markedly suppressed. At 1 s*^−^*^1^, only the highest strain magnitude (0.5) produced a modest but significant decrease in mean event rate when comparing 0+ vs 0- (Fig. 2B; mean ΔER = *−*24.24%, 95% CI [-41.70, -1.55]%; *N* = 6; *z* = *−*2.08, *p* = 0.038, Tweedie GLMM). Thus, increasing strain alone weakly suppresses activity at slow loading, whereas rapid loading produces a substantially greater suppression of activity, indicating a strain-rate–dependent transition.

**Figure 2.**
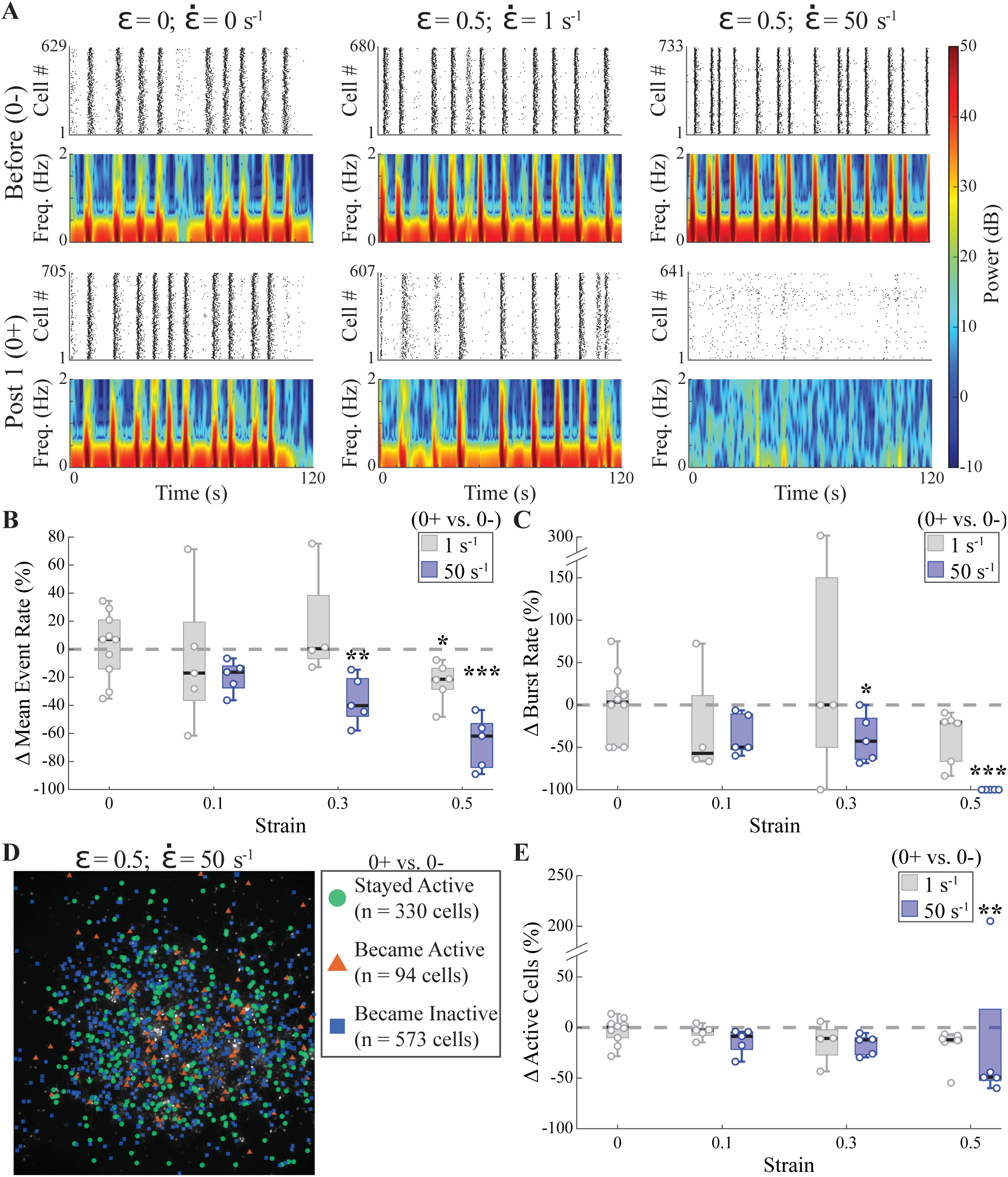
Strain rate sets the threshold for hyperacute network dysfunction. (**A**) Representative rasters and spectrograms before (0-) and after (0+) stretch; the largest disruptions occur at higher strain rate. (**B**) Change in mean event rate among active cells (0+ vs 0-) across strains and rates (box plots; *N* = 4–10 cultures per condition). (**C**) Change in network burst rate (0+ vs 0-) across strains and rates (box plots; *N* = 4 *−* 10 cultures per condition); significant decreases at *ε* = 0.3 and 0.5, 50 s*^−^*^1^. No culture stretched to *ε* = 0.5 at 50 s*^−^*^1^ had bursts at 0+; significance for that condition reflects a two-sided Fisher’s exact test of the proportion of cultures without bursts against sham. (**D**) Spatial classification for a representative *ε* = 0.5, 50 s*^−^*^1^ field showing cells that stayed active, became active, or became inactive from 0- to 0+. (**E**) Change in number of active cells (0+ vs 0-) across strains and rates (box plots; *N* = 4–10 cultures per condition). Statistics: \**p <* 0.05, \*\**p <* 0.01, \*\*\**p <* 0.001. *p* = 0.0053, Tweedie GLMM). This reduction was directionally consistent across samples but borderline at the 0.05 threshold under leave-one-out sensitivity analysis (Table S1).

We next asked whether this activity suppression altered network synchrony, quantified by burst rate (bursts/min). Bursts, defined as *≥* 20% active cells within a continuous time window, are a sign of a healthy connected network *in vitro*. In the representative sample stretched at 50 s*^−^*^1^, network bursts are effectively eliminated, although isolated synchronous attempts remain visible in the raster (Fig. 2A). This loss of synchrony is confirmed at the population level (Fig. 2C). Only the 0.3 and 0.5 strains at 50 s*^−^*^1^ showed significantly altered burst rates. There was an average 44% decrease in burst rate within the first two-minutes of stretch at 0.3 strain (Fig. 2C; mean Δburst rate = *−*44.14%, 95% CI [-64.54, -12.01]%; *N* = 5; *z* = *−*2.51, *p* = 0.012, Tweedie GLMM). In all five samples stretched to 0.5 strain at 50 s*^−^*^1^, network bursts were completely abolished at 0+ (Fig. 2C; mean Δburst rate = *−*100%; *N* = 5). Because every culture in this condition reached zero, the outcome is completely separated and no Wald test is interpretable; comparing the proportion of cultures without bursts at 0+ against sham (5 of 5 versus 0 of 10) gives *p* = 3.3 *×* 10*^−^*^4^, Fisher’s exact test. This indicates near-total decoupling of network synchrony.

Finally, we tested whether neurons ceased participating in activity altogether by classifying cells as active (*≥* 1 event) or inactive (Fig. 2D). Only 0.5 strain at 50 s*^−^*^1^ significantly reduced the fraction of active neurons (Fig. 2E; 0.5 @ 50 s*^−^*^1^ mean Δactive cells = *−*29.43%, 95% CI [-44.75, -9.86]%; *N* = 5; *z* = *−*2.79,

### Rapid loading reveals a threshold for calcium homeostasis disruption

The previous section demonstrated that rapid loading suppresses calcium activity at 0+. We next asked whether this suppression reflects only altered spiking dynamics or whether a distinct change in calcium handling emerges at higher mechanical loading. Because traumatic injury is often accompanied by elevations in intracellular calcium (33, 52–56), we evaluated shifts in basal fluorescence as an indicator of altered calcium homeostasis. Here, ‘basal fluorescence’ refers to the steady-state GCaMP signal outside of transient events. Although GCaMP does not report absolute calcium concentration, sustained changes in basal fluorescence indicate a shift in intracellular calcium set-point within the measurable dynamic range.

To distinguish measurement variability and photobleaching from persistent cellular fluxes, a threshold was applied to changes in basal fluorescence (see Methods). Figure 3A-B illustrates the cellular classifications in representative sham and 0.5 strain @ 50 s*^−^*^1^ samples. Insets adjacent to each image show the raw basal fluorescence of a zoomed-in section at 0- and 0+. The sham sample shows minimal basal change (Fig. 3A). In contrast, the 0.5 strain at 50 s*^−^*^1^ condition exhibits a large population of cells with elevated basal fluorescence (Fig. 3B). This network-level increase is further illustrated by the distributional differences between the representative samples in Fig. 3C. When comparing strain and strain rate effects across conditions, a significant increase in the percentage of cells with elevated basal fluorescence occurs only at 0.5 strain @ 50 s*^−^*^1^ (Fig. 3D; estimated marginal mean = 42.60% of neurons, 95% CI [28.88, 56.32]%, versus 4.69% [1.63, 7.74]% in sham; difference = +37.92 percentage points, 95% CI [19.42, 56.41]; *N* = 5; *z* = 5.29, *p* = 7.3 *×* 10*^−^*^7^, Dunnett-adjusted beta-binomial model). Because elevated basal fluorescence could reduce transient detectability within the indicator’s dynamic range, we explicitly evaluated signal saturation in this condition (Supporting Material, Fig. S3). These analyses confirm that transient suppression reflects altered dynamics rather than measurement ceiling effects. All other conditions show no significant basal fluorescence shift, indicating that altered calcium homeostasis emerges only beyond a mechanical loading threshold.

**Figure 3.**
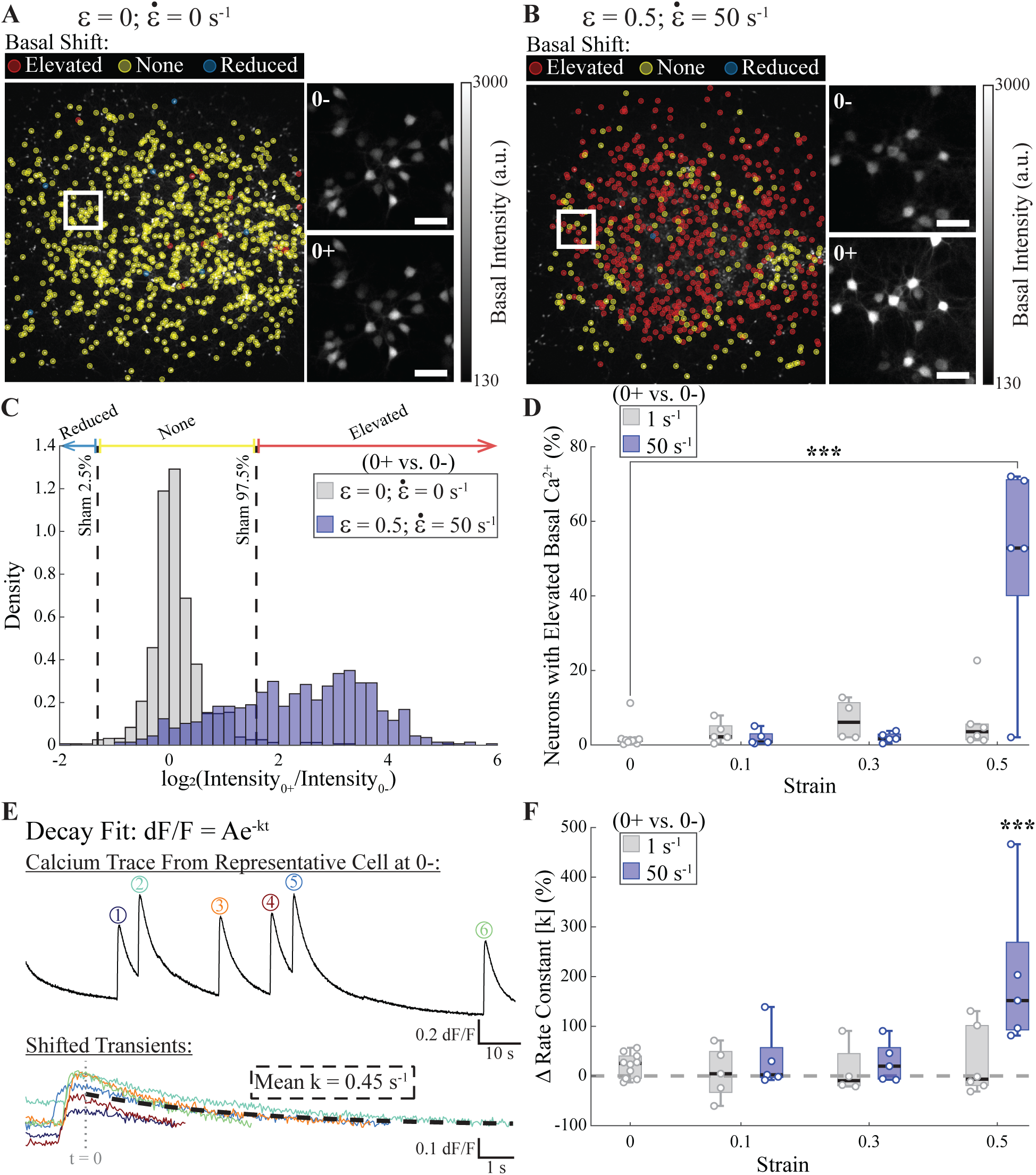
Calcium homeostasis distorts immediately after high strain and strain rate loading. (**A-B**) Spatial classification of basal GCaMP fluorescence intensity shifts (0+ vs 0-) for representative sham and *ε* = 0.5 @ 50 s*^−^*^1^ fields. Insets sho w raw basal fluorescenc e images used for the shift calculation (scale bar, 50 *µ*m). (**C**) Probability densities of log_2_ Intensity_0+_*/*Intensity_0_*_−_* for samples in A-B; vertical dashed lines mark sham-derived 95th-percentile thresholds for meaningful reduction/elevation. (**D**) Percentage of neurons with elevated basal fluorescence (0+ vs 0-) across strains and rates (box plots; *N* = 4 *−* 10 cultures per condition). (**E**) Representative calcium transient with exponential fit used to compute the rate constant, *k*, of the decay (black dashed line). (**F**) Change in mean rate constant (0+ vs 0-) across strains and rates (box plots; *N* = 4 *−* 10 cultures per condition). Statistics: \*\*\**p <* 0.001.

We next tested whether calcium clearance dynamics also change at this loading threshold. In GCaMP6s, the calcium transient decay rate reflects processes that remove or buffer cytosolic calcium following influx (57, 58). Decay rate constants were obtained by fitting an exponential to individual calcium transients (Fig. 3E; described in the Methods). As with the basal fluorescence shift, the only significant change in the average transient decay rate constant (*k*) occurs at 0.5 strain @ 50 s*^−^*^1^ (Fig. 3F; mean Δ*k* = 186.46%, 95% CI [108.14, 294.23]%; *N* = 5; *z* = 6.46, *p* = 1.05 *×* 10*^−^*^10^, Gamma GLMM). Together, elevated basal fluorescence and accelerated decay indicate the emergence of an altered calcium homeostatic state rather than a change in calcium dynamics alone. The accelerated decay may reflect increased engagement of calcium clearance mechanisms, such as ATP-dependent pumps (e.g., SERCA or PMCA), in response to elevated intracellular calcium levels.

### Neuronal activity partially recovers within 24 hours despite reduced neuronal participation

Next, we asked whether the hyperacute network dysfunction resolved or evolved into an acute change in network activity by analyzing the same samples 24 hours later. Figure 4A compares the single-cell event rate distributions across time for representative sham and *ε* = 0.5 @ 50 s*^−^*^1^ samples. The stretch condition shows a redistribution of event rates among active neurons that is absent in sham cultures. When examining recovery of average activity (24h vs 0+), most conditions show no significant change (Fig. 4B). The condition *ε* = 0.1 @ 1 s*^−^*^1^ shows a trend toward decreased mean event rate (Fig. 4B; mean ΔER between *−*54% and *−*58%; *N* = 5, Tweedie GLMM) that was not robust to model specification (Šidák-adjusted *p* = 0.016–0.059 across model fits). Conversely, *ε* = 0.5 @ 50 s*^−^*^1^ shows a significantly increased mean event rate (Fig. 4B; mean ΔER = 163.05%, 95% CI [39.53, 395.92]%; *N* = 5; *z* = 3.41, *p* = 0.0013, Tweedie GLMM). Despite this apparent increase in mean event rate, network burst activity did not recover by 24h (Fig. S4B), although bursting was also absent at 24h in 5 of 10 sham cultures (Supporting Material). So, the remaining active cells did not restore coordinated network bursting.

**Figure 4.**
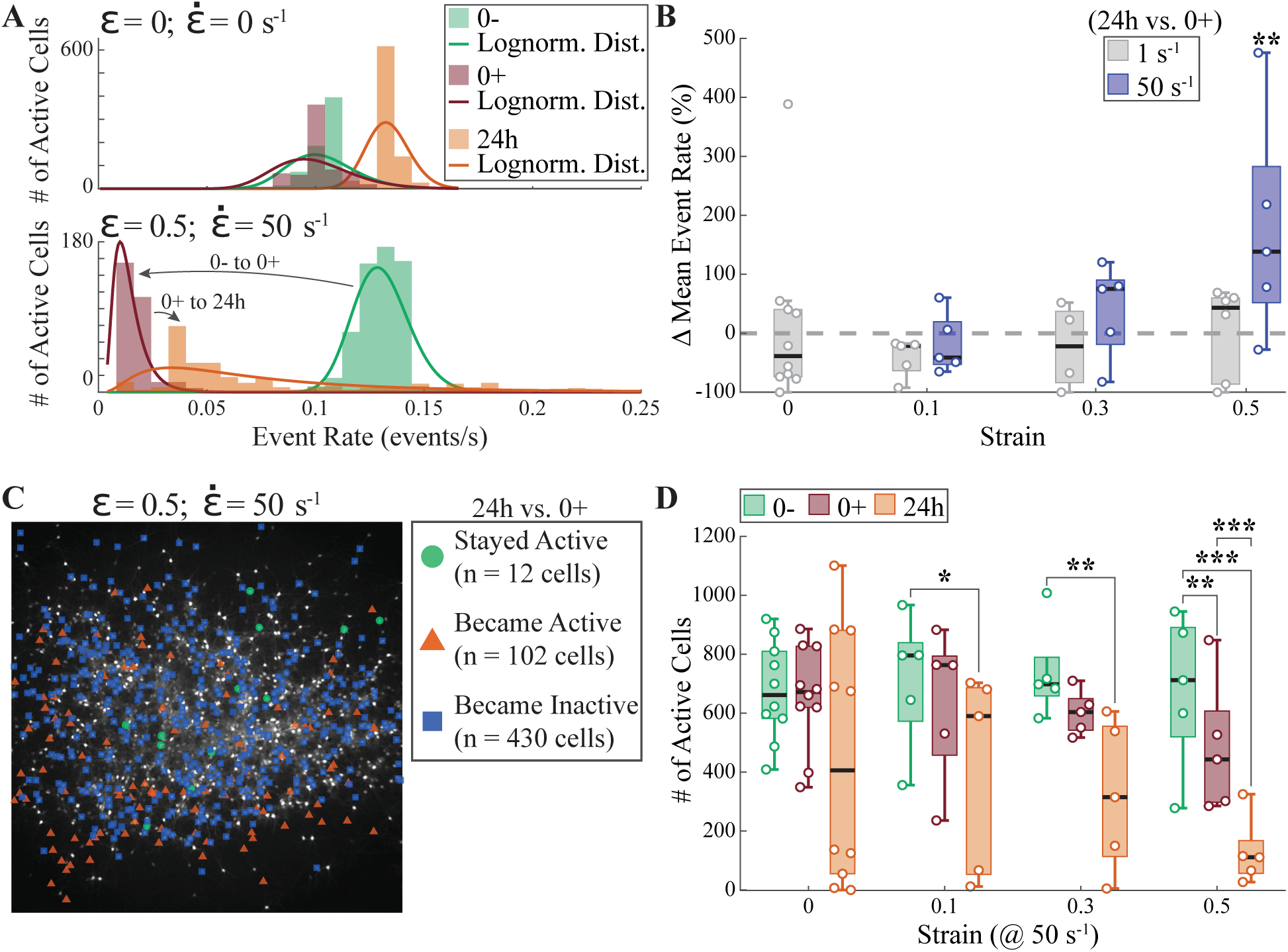
Network attrition occurs over 24 hours despite recovery in remaining active cells in severe loading conditions. (**A**) Histograms showing shifts in per-cell event rate distribution between 0-, 0+, and 24h for representative sham and 0.5 strain at 50 s*^−^*^1^ samples. (**B**) Change in mean event rate among active cells (24h vs 0+) across strains and strain rates (box plots of per-sample percent change; *N* = 4 *−* 10 cultures per condition). (**C**) Spatial classification for a representative *ε* = 0.5, 50 s*^−^*^1^ field showing cells that stayed active, became active, or became inactive from 0+ to 24h. (**D**) Number of active cells at 0-, 0+, and 24h across strains at 50 s*^−^*^1^ (box plots; *N* = 4 *−* 10 cultures per condition). Statistics: \**p <* 0.05, \*\**p <* 0.01, \*\*\**p <* 0.001.

To determine whether this apparent recovery reflected restoration of the original network or changes in neuronal participation, we quantified the number of active cells. In the most severe stretch condition of *ε* = 0.5 @ 50 s*^−^*^1^, the number of active cells decreases despite the increased mean event rate among those remaining active (Fig. 4C). Within the 50 s*^−^*^1^ condition, neither *ε* = 0.1 nor *ε* = 0.3 shows a significant change in the number of active cells immediately after stretch (0+ vs 0-; *ε* = 0.1: mean Δ = *−*10.79%, 95% CI [-28.73, 11.65]%, *N* = 5, *z* = *−*1.00, *p* = 0.319; *ε* = 0.3: mean Δ = *−*18.40%, 95% CI [-34.89, 2.27]%, *N* = 5, *z* = *−*1.76, *p* = 0.078; Tweedie GLMM). However, both conditions exhibit significant reductions in the number of active cells by 24h (Fig. 4D). Relative to baseline (0-), active cells decreased by 45.13% at *ε* = 0.1 @ 50 s*^−^*^1^ (95% CI [-67.37, -7.72]%; *N* = 5; *z* = *−*2.58, *p* = 0.020, Tweedie GLMM) and by 56.23% at *ε* = 0.3 @ 50 s*^−^*^1^ (95% CI [-76.52, -18.41]%; *N* = 5; *z* = *−*2.97, *p* = 0.006, Tweedie GLMM). These decreases were directionally consistent across samples; the *ε* = 0.3 @ 50 s*^−^*^1^ result was borderline at the threshold under leave-one-out sensitivity analysis (Table S1). Sham cultures, however, also lost active cells over 24 hours, although less consistently (mean Δ = *−*33.25%, 95% CI [-63.06, 20.59]%; *N* = 10; *z* = *−*1.53, *p* = 0.237, Tweedie GLMM), and neither decrease differed significantly from the change in sham (*ε* = 0.1: *p* = 0.98; *ε* = 0.3: *p* = 0.68; Dunnett-adjusted comparison of the 24h vs 0- change against sham). For *ε* = 0.5 @ 50 s*^−^*^1^, there was a significant decrease in the number of active cells between 0+ vs 0- and an even larger decrease between 24h vs 0- (Fig. 4D; 24h vs 0-: mean Δactive cells = *−*81.10%, 95% CI [-91.14, -59.70]%, *z* = *−*4.92, *p* = 1.73 *×* 10*^−^*^6^; 24h vs 0+: mean Δactive cells = *−*73.22%, 95% CI [-87.74, -41.53]%, *z* = *−*3.77, *p* = 3.22 *×* 10*^−^*^4^; all *N* = 5, Tweedie GLMM); this 24h decrease also exceeded the change in sham (ratio of changes = 0.28, 95% CI [0.09, 0.86]; *z* = *−*2.93, *p* = 0.018). Changes in cell participation across strains at the 1 s*^−^*^1^ condition are shown in Fig. S4C.

Thus, the recovery of average neuronal activity arises from increased activity within a smaller subset of neurons rather than restoration of the original network activity pattern. The 24-hour calcium dynamic response reflects partial recovery of activity levels alongside reduced cellular participation, indicating a persistent shift in overall network activity. Spatial analysis of pairwise functional connectivity independently confirmed this pattern: the fraction of connected cell pairs collapsed acutely in *ε* = 0.5 @ 50 s*^−^*^1^ cultures (*p* = 0.004, Tukey-adjusted LME) and remained suppressed at 24 hours (*p* = 0.002), while Moran’s *I* spatial autocorrelation was not significantly altered in any condition (all *p >* 0.82), indicating global rather than spatially patterned connectivity loss (Fig. S5). This pattern may reflect compensatory hyperactivity in a subset of resilient neurons, potentially driven by disinhibition or altered calcium homeostasis following injury.

### Stretch-induced functional changes occur without loss of cell viability

Our analyses thus far show that cultures stretched to 0.5 strain at 50 s*^−^*^1^ exhibit reduced neuronal participation in network activity 24 hours after injury, beyond the change observed in sham cultures. To determine whether this reduction reflects functional changes in network activity rather than injury-induced cell loss, we measured cell viability in the same cultures used for calcium imaging. Hoechst staining was used to label nuclei of viable cells, while Ethidium homodimer labeling identified cells with compromised membranes indicative of cell death.

Cells were classified based on their Hoechst and Ethidium signals across time points. Hoechst staining labels nuclei of viable cells, whereas Ethidium homodimer enters cells with compromised membranes. Cells were therefore categorized into three groups: (1) surviving cells that retained Hoechst signal and were Ethidium-negative, (2) cells that lost Hoechst signal between 0- and 24h, indicating nuclear degradation following cell death, and (3) cells that remained Hoechst-positive but became Ethidium-positive at 24h, indicating membrane compromise. Cells in categories (2) and (3) were combined and quantified as dead cells (Fig. 5A-C). Detailed classification criteria are described in the Methods.

**Figure 5.**
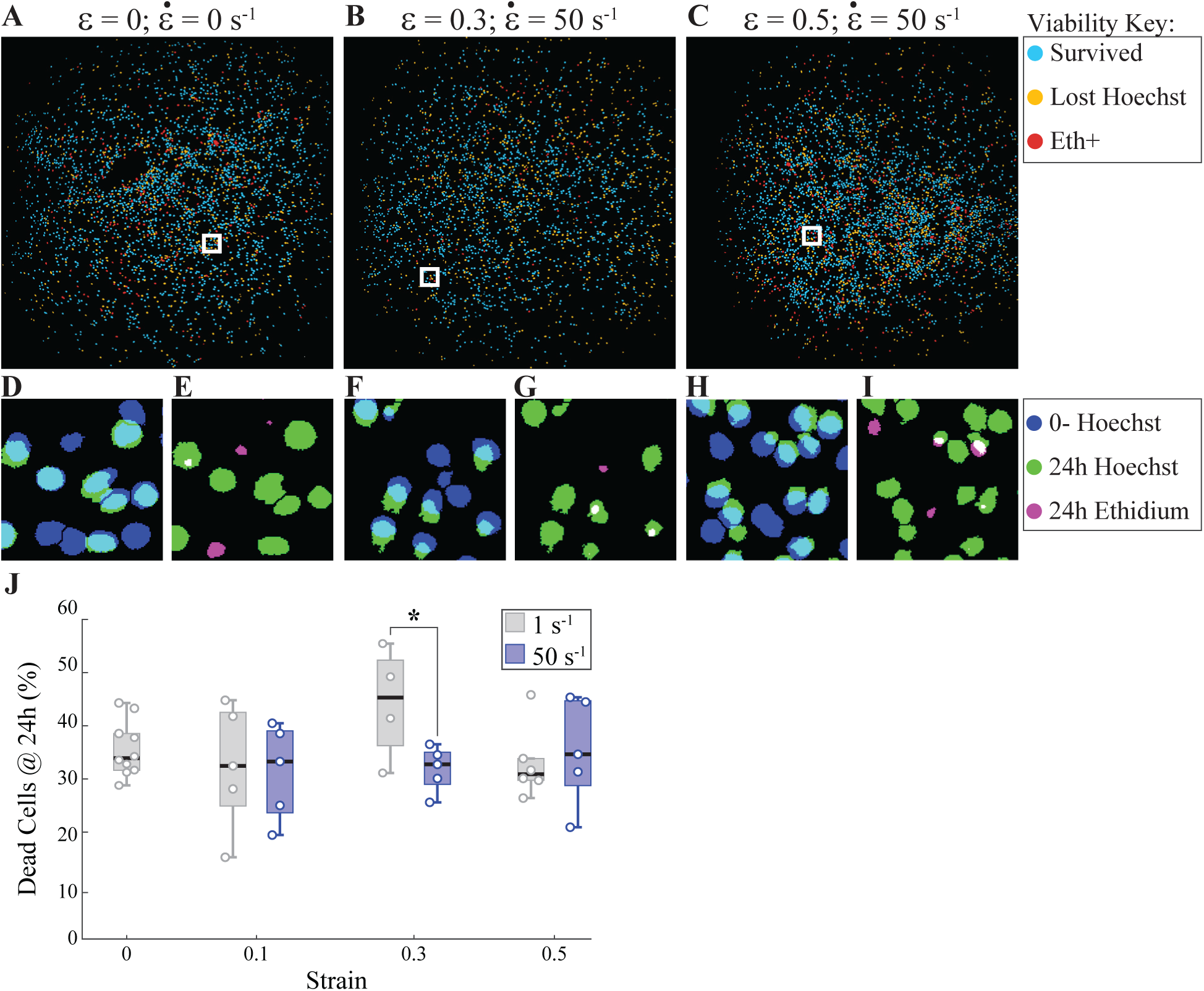
Cell viability at 24 hours following stretch injury. (**A–C**) Per-nucleus classification overlays for representative sham, 0.3 @ 50 s*^−^*^1^, and 0.5 @ 50 s*^−^*^1^ fields showing survived cells (light blue), nuclei that lost Hoechst signal between 0- and 24h (yellow), and Ethidium-positive (Eth+) nuclei at 24h (red). White boxes indicate the regions shown in (D–I) that were chosen to exemplify the viability classification. (**D,F,H**) Hoechst signal at 0- (blue) and 24h (green), illustrating nuclei retained versus lost at 24h. Minor non-overlap within a conservative drift threshold was classified as survived (see Methods). (**E,G,I**) Hoechst at 24h (green) and Ethidium at 24h (magenta), highlighting Eth+ nuclei. (**J**) Population-level quantification of percent dead cells relative to 0- Hoechst counts across strain magnitudes and strain rates (Dead Cells = [cells*_Lost_*__*Hoechst*_ + cells*_Eth_*_+_]*/*cells_0_*_−_ ∗* 100; *N* = 4 *−* 10 cultures per condition). \**p <* 0.05.

Representative samples show no qualitative differences between sham cultures and samples stretched at 50 s*^−^*^1^. Example regions illustrate comparable patterns of nuclear loss and membrane compromise across conditions (Fig. 5D–I). At the population level, the percentage of dead cells does not differ significantly between sham samples and any stretch condition tested (Fig. 5J; all sham vs stretch comparisons *p >* 0.26, Dunnett-adjusted beta-binomial model). The only statistically significant difference occurs between strain-rate conditions at *ε* = 0.3, where the 50 s*^−^*^1^ condition shows a modest reduction in cell death relative to 1 s*^−^*^1^ (difference = *−*12.01 percentage points, 95% CI [-21.96, -2.06]; *z* = *−*2.37, *p* = 0.018, beta-binomial model, planned unadjusted contrast). This reduction was directionally consistent across samples but borderline at the 0.05 threshold under leave-one-out sensitivity analysis (Table S1).

Thus, although rapid loading produces substantial changes in neuronal activity and cellular participation, these functional alterations occur without detectable loss of cell viability within 24 hours of injury. Together with the observed shifts in calcium homeostasis, these findings indicate that mechanical stretch induces persistent functional dysregulation in surviving neurons rather than widespread neuronal death.

## Discussion

We used high-precision, controlled uniaxial tension that mimics tensile deformation in mTBI at the cellular scale and combined this system with high-speed imaging of neuronal network calcium dynamics. This approach resolves the hyperacute neuronal response to mechanical stretch and shows that deformation rapidly disrupts network activity in a loading-dependent manner. The experiments presented here reveal that mechanical deformation induces strain- and strain-rate-dependent modes of network disruption within seconds of injury, ranging from graded hypoactivity to abrupt collapse of synchronous bursting and concurrent calcium homeostasis failure at distinct mechanical thresholds. Our analysis places secondary injury within a time-dependent process, highlighting cellular dynamics that have remained ambiguous across TBI phases (8, 59). Most experimental paradigms have detected functional abnormalities only minutes to hours after insult, thereby excluding the hyperacute phase (30, 31). Single-cell-resolved analysis immediately following stretch establishes that the earliest seconds after deformation shape not only the immediate signaling phenotype but also the trajectory of network recovery over the subsequent 24 hours, providing a framework that separates the primary mechanical insult from secondary functional pathology in mTBI. These disruptions provide a potential functional basis for the cognitive and memory impairments observed following traumatic brain injury (60) and may also be relevant to hyperacute clinical phenomena such as brief loss of consciousness (23), suggesting that such deficits may originate from an immediate breakdown of neuronal communication rather than from delayed structural damage.

We observe that mechanical deformation initiates secondary cascades that rapidly impair neuronal network function without necessarily producing overt structural or morphological phenotypes. We demonstrate that hyperacute signaling disruptions bifurcate into distinct modes governed by strain and strain-rate thresholds. Rapid loading rates (50 s*^−^*^1^) trigger graded attenuation beginning at moderate strain (0.3), with disruption escalating at higher strain. At a strain of 0.5, we observe complete collapse of network bursting together with calcium homeostasis failure, reflected by elevated basal levels and accelerated transient decay. These thresholds demarcate the isolated loss of excitability from ionic dysregulation, which progresses to network decoupling, and suggest that mechanical parameters dictate not only injury severity but also the nuanced biophysical mode of cellular response. The frequency of calcium transients depended on both strain magnitude and strain rate: high-rate loading reduced event frequency at lower strains than low-rate stretch, and the degree of dysfunction scaled with strain under high-rate conditions. High-rate loading likely disrupts cellular signaling more abruptly, whereas lower-rate stretch permits partial adaptation. Network-level analysis further clarified the nature of suppression in the most severe condition. Specifically, a strain of 0.5 applied at 50 s*^−^*^1^ abolished synchronous network bursts across all cultures. Because network bursts reflect coordinated synaptic communication rather than isolated firing, their disappearance indicates a potential breakdown of functional connectivity rather than a simple reduction in excitability. This aligns well with previous work on mild TBI in rodents, demonstrating reduction in low frequency calcium transients together with frequency dependent changes in global functional connectivity (61). Future work using our system could unravel these functional connectivity changes in the hyperacute stage. Thus, moderate stretch primarily suppresses participation in calcium-event activity, whereas threshold-level stretch also collapses coordinated network communication, triggering an additional mode of network failure. This hyperacute response aligns with biomechanical studies showing that cellular injury depends strongly on strain rate and that cells can often accommodate slower loading more effectively than rapid deformation (62). Consistent with this principle, neuronal injury studies report greater calcium dysregulation, synaptic dysfunction, and pathology at higher strain rates (27, 54). The abrupt elimination of synchronous bursts, a hallmark of coordinated synaptic transmission, indicates functional disconnection rather than uniform silencing and provides a potential basis for the acute connectivity deficits observed clinically after mTBI by fMRI and EEG (63–65).

These findings are consistent with recruitment of parallel mechanosensitive channels rather than a single monotonic calcium response. The early reduction in calcium-transient activity at lower loading severities aligns with the activation of stretch-sensitive TREK-1/TRAAK channels, which dampen neuronal excitability during axonal stretch (39). By contrast, the basal fluorescence elevation that emerged only in the most severe rapid-loading condition corresponds more with mechanically triggered calcium-entry pathways such as NMDA receptor and TRPV4 signaling (35–37). Neuronal TRPV4 can mediate mechanically induced calcium and sodium influx (37), whereas astrocytic Piezo1 and TRPV4 signaling can evoke calcium responses together with ATP or gliotransmitter release (38, 66). Together, these pathways provide plausible links between severe stretch, neuronal calcium dyshomeostasis, and altered network communication in our mixed cultures. In this framework, reduced calcium transients and elevated basal fluorescence need not originate from the same channel in the same cell, which is congruent with our single-cell finding that these two hyperacute phenotypes were not tightly coupled.

In line with this mechanistic backdrop, the present neuronal measurements revealed two co-occurring yet distinct components of injury: an immediate decrease in calcium transients and a disturbance in ionic homeostasis that emerged only beyond a mechanical threshold. Importantly, at 0.5 strain and 50 s*^−^*^1^, cytosolic calcium increased, as reflected by elevated basal fluorescence, while transient activity remained diminished. This indicates that severe loading does not replace hypoactivity with calcium dysregulation, but rather superimposes ionic dysregulation onto an already depressed activity state. Single-cell analysis further showed no clear association between reduced activity and elevated basal fluorescence (Fig. S6), signifying that calcium overload was not restricted to the same cells that lost calcium-event activity. This pattern is consistent with prior work showing that mechanical injury induces rapid neuronal depolarization through disruption of ionic homeostasis and neurotransmitter release (8, 67). Instead of increasing firing rate, sustained depolarization can inactivate voltage-gated sodium channels and suppress spike generation (68). This phenomenon is observed during the early stages of spreading depolarization, where the local site of insult undergoes immediate membrane depolarization together with suppression of high-frequency activity (67, 69). Because somatic calcium transients are tightly coupled to action potential firing (70, 71), this mechanism provides a plausible explanation for the immediate reduction in calcium events observed here. In parallel, the elevation in basal fluorescence under high-rate, high-magnitude stretch is consistent with the rapid calcium influx documented immediately after TBI (33, 52–56). Given the seconds-scale onset observed here, activation of NMDA receptors is a plausible contributor because glutamate-mediated calcium entry occurs rapidly after mechanical perturbation (33, 72). Once intracellular calcium reaches a threshold, additional influx through voltage-gated calcium channels and release from intracellular stores may further amplify the response (72). Together, these findings indicate that severe loading produces a combined state of reduced spike-coupled calcium-event activity and elevated cytosolic calcium, consistent with a spreading-depolarization-like response in which depolarization-associated loss of excitability occurs alongside calcium influx through mechanosensitive and downstream voltage-gated pathways (30, 56, 73, 74). Additionally, the accelerated decay kinetics observed under this same severe condition also suggest recruitment of calcium-clearance mechanisms, potentially including ATP-dependent pumps such as PMCAs, in response to elevated cytosolic calcium (75). More broadly, this pattern supports calcium dysregulation and calcium-handling pathways as potential therapeutic targets for limiting downstream dysfunction after mild traumatic brain injury (53, 76).

At 24 hours after the most severe loading (0.5 strain at 50 s*^−^*^1^), event rates partially recover in the remaining active neurons, yet overall participation remains depressed beyond the change observed in sham cultures. This selective attrition, together with compensatory hyperactivity in persistently active cells, implies network reorganization rather than merely transient suppression. Importantly, reduced neuronal participation at 24 hours was not attributable to cell death, consistent with prior *in vitro* stretch models showing minimal cell death under similar conditions (28, 31, 33, 34, 77, 78). Even the most severe stretch profile applied here, which triggered a clear disorder in basal cytosolic calcium, was insufficient to induce significant cell death by 24 hours. This supports the interpretation that the applied loading conditions model mTBI, which is characterized by functional impairment without widespread degeneration. The observed pattern is also consistent with emerging mild stretch models showing sublethal calcium and metabolic dysfunction (79, 80), reinforcing the idea that under mild-to-moderate loading, injury may be dominated by circuit-level dysfunction rather than necrotic loss.

While these interpretations are supported by the observed relationships, several limitations define the scope of this study and inform future work. All cultures were derived from female pups. Sex differences have been reported in *in vitro* traumatic axonal injury (81), though whether dissociated postnatal cultures retain them is unresolved; the single-sex design was intended to remove that variability rather than test it. Sham cultures also lost activity over 24 hours, most markedly in cultures prepared between July and September; 24-hour changes in participation are therefore interpreted relative to sham, against which only the most severe loading condition differed. The present imaging approach was optimized for hyperacute, large-field, single-cell tracking of calcium activity across extensive neuronal populations rather than for resolving calcium dynamics within specific intracellular compartments or quantifying absolute calcium concentration. Complementary approaches, including ratiometric calcium indicators and organelle-targeted measurements of mitochondrial or endoplasmic reticulum calcium, would help refine the magnitude, source, and persistence of post-injury calcium elevations (82–84). Notably, related stretch-injury studies have already used Fura-2 to examine post-traumatic calcium handling and intracellular store signaling, indicating that these methods would extend the present network-level framework (85). An additional consideration for interpreting altered decay kinetics and basal fluorescence shifts is that GCaMP behavior under acute mechanical deformation has not been directly characterized. Although a fraction of cellular CaM — estimated at approximately 15% in brain tissue — is membrane-associated (86), the majority of cytoplasmic GCaMP6s lacks a defined mechanical load path (87–89), making the fluorescence signal unlikely to be substantially affected by direct mechanical action. Our system offers a distilled platform for answering questions of rapid effects in mTBI. Capturing the full cellular and structural complexity of brain tissue will, however, require the development of a new set of stretch devices that mimic injury propagation and recovery. Future studies using such technologies could likely integrate population-scale volumetric functional imaging with complementary electrophysiological and quantitative calcium approaches in more complex models and will be important for defining longer-term recovery and the molecular basis of the distinct injury modes identified here.

## Conclusion

Collectively, these results refine insights gained from purely graded injury models by establishing that hyperacute mechanical signatures encode downstream cascade selection. Early strain-rate thresholds may therefore serve as predictive determinants of secondary progression, helping explain variability in mTBI outcomes and the limitations of diagnoses based solely on later measurements. Targeting these earliest modes of network failure offers a precise window for intervention before compensatory remodeling entrenches chronic dysfunction. This framework advances mechanistic understanding of mTBI pathophysiology and supports the rational design of injury-specific preclinical models.

## Data and code availability

Processed datasets, supplemental summary data, intermediate cell-viability masks, and sample metadata supporting the findings of this study are available in the Dryad Digital Repository at https://doi.org/10.5061/dryad.1jwstqk9h (90). Raw calcium-imaging movies are available from the corresponding author on reasonable request owing to their large file size. All original analysis code (cell-viability, spatial network, and statistical modeling routines in MATLAB and R) is freely available in the same Dryad deposit and is publicly accessible as of the date of publication. Calcium signal detection and characterization were performed using NeuroSpikeX (v1.0.0) (51), freely available at https://github.com/FranckLab/NeuroSpikeX. Soma detection and fluorescence extraction used NeuroCa (50).

## Author Contributions

Jamie A. Sergay: Methodology, Software, Investigation, Formal analysis, Data curation, Visualization, Writing – original draft, Writing – review & editing. Natalie Schick: Formal analysis, Writing – review & editing. Emily Blick: Investigation, Writing – review & editing. Mauricio Araiza Canizales: Formal analysis, Visualization. Jing Zhang: Conceptualization, Resources, Project administration. Christian Franck: Conceptualization, Funding acquisition, Supervision, Writing – review & editing. Aviad Hai: Conceptualization, Funding acquisition, Supervision, Writing – review & editing.

## Declaration of generative AI and AI-assisted technologies in the writing process

The authors used OpenAI ChatGPT (GPT-4o) to assist with developing, refining, and troubleshooting custom MATLAB and R code used for calcium-signal analyses, cell-viability image analysis, supplementary analyses, and statistical modeling. ChatGPT was also used to support language editing, organization, and clarity during manuscript preparation. All AI-assisted code and text were reviewed, edited, and verified by the authors, who take full responsibility for the final manuscript, analyses, interpretations, and conclusions. No generative AI-created images, schematics, multimedia, primary data, or citations were included in this manuscript.

## Acknowledgments

We would like to thank Dr. Kip Ludwig for his feedback and advice. This material is based upon research supported by the U.S. Office of Naval Research under PANTHER award numbers N00014-22-1-2828 and N00014-23-1-2006 through Dr. Timothy Bentley.

## Supporting Material

Supporting Material, provided as a separate file, includes supporting methods, supporting text, six figures (Figs. S1–S6), and one table (Table S1).

## References

1. DeKosky, S. T., and B. M. Asken, 2017. Injury cascades in TBI-related neurodegeneration. Brain Inj. 31:1177–1182.

2. Ng, S. Y., and A. Y. W. Lee, 2019. Traumatic brain injuries: Pathophysiology and potential therapeutic targets. Front. Cell. Neurosci. 13:528.

3. Curley, W. H., Y. G. Bodien, D. W. Zhou, M. M. Conte, A. S. Foulkes, J. T. Giacino, J. D. Victor, N. D. Schiff, and B. L. Edlow, 2022. Electrophysiological correlates of thalamocortical function in acute severe traumatic brain injury. Cortex 152:136–152.

4. Verboom, M., R. van den Berg, M. van de Ruit, and M. van der Jagt, 2026. Prognostic value of electroencephalography in critically ill adult patients with traumatic brain injury: A systematic review. J. Neurotrauma 43:13–34.

5. Leskinen, S., N. H. Mehta, H. A. Shah, M. Quelle, R. Woodworth, G. Dituri, C. Balucani, and R. S. D’Amico, 2025. Functional connectivity changes in traumatic brain injury: A systematic review and coordinate-based meta-analysis of fMRI studies: A systematic review and coordinate-based meta-analysis of fMRI studies. Neurology 105:e214298.

6. Demirtas-Tatlidede, A., A. M. Vahabzadeh-Hagh, M. Bernabeu, J. M. Tormos, and A. Pascual-Leone, 2012. Noninvasive brain stimulation in traumatic brain injury. J. Head Trauma Rehabil. 27:274–292.

7. Dewan, M. C., A. Rattani, S. Gupta, R. E. Baticulon, Y.-C. Hung, M. Punchak, A. Agrawal, A. O. Adeleye, M. G. Shrime, A. M. Rubiano, J. V. Rosenfeld, and K. B. Park, 2018. Estimating the global incidence of traumatic brain injury. J. Neurosurg. 1–18.

8. MacFarlane, M. P., and T. C. Glenn, 2015. Neurochemical cascade of concussion. Brain Inj. 29:139–153.

9. Guerriero, R. M., C. C. Giza, and A. Rotenberg, 2015. Glutamate and GABA imbalance following traumatic brain injury. Curr. Neurol. Neurosci. Rep. 15:27.

10. Yasen, A. L., J. Smith, and A. D. Christie, 2018. Glutamate and GABA concentrations following mild traumatic brain injury: a pilot study. J. Neurophysiol. 120:1318–1322.

11. Vascak, M., X. Jin, K. M. Jacobs, and J. T. Povlishock, 2018. Mild traumatic brain injury induces structural and functional disconnection of local neocortical inhibitory networks via parvalbumin interneuron diffuse axonal injury. Cereb. Cortex 28:1625–1644.

12. McKee, A. C., and M. E. Robinson, 2014. Military-related traumatic brain injury and neurodegeneration. Alzheimers. Dement. 10:S242–53.

13. Zhou, Y., M. P. Milham, Y. W. Lui, L. Miles, J. Reaume, D. K. Sodickson, R. I. Grossman, and Y. Ge, 2012. Default-mode network disruption in mild traumatic brain injury. Radiology 265:882–892.

14. Iraji, A., R. R. Benson, R. D. Welch, B. J. O’Neil, J. L. Woodard, S. I. Ayaz, A. Kulek, V. Mika, P. Medado, H. Soltanian-Zadeh, T. Liu, E. M. Haacke, and Z. Kou, 2015. Resting state functional connectivity in mild traumatic brain injury at the acute stage: Independent component and seed-based analyses. J. Neurotrauma 32:1031–1045.

15. Kang, W. H., and B. Morrison, 3rd, 2015. Predicting changes in cortical electrophysiological function after in vitro traumatic brain injury. Biomech. Model. Mechanobiol. 14:1033–1044.

16. Dimitriadis, S. I., G. Zouridakis, R. Rezaie, A. Babajani-Feremi, and A. C. Papanicolaou, 2015. Functional connectivity changes detected with magnetoencephalography after mild traumatic brain injury. NeuroImage Clin. 9:519–531.

17. Zhang, J., K. Safar, Z. Emami, G. M. Ibrahim, S. E. Scratch, L. da Costa, and B. T. Dunkley, 2020. Local and large-scale beta oscillatory dysfunction in males with mild traumatic brain injury. J. Neurophysiol. 124:1948–1958.

18. Dall’Acqua, P., S. Johannes, L. Mica, H.-P. Simmen, R. Glaab, J. Fandino, M. Schwendinger, C. Meier, E. J. Ulbrich, A. Müller, H. Baetschmann, L. Jäncke, and J. Hänggi, 2017. Functional and structural network recovery after mild traumatic brain injury: A 1-year longitudinal study. Front. Hum. Neurosci. 11:280.

19. Holden, S. S., F. C. Grandi, O. Aboubakr, B. Higashikubo, F. S. Cho, A. H. Chang, A. O. Forero, A. R. Morningstar, V. Mathur, L. J. Kuhn, P. Suri, S. Sankaranarayanan, Y. Andrews-Zwilling, A. J. Tenner, A. Luthi, E. Aronica, M. R. Corces, T. Yednock, and J. T. Paz, 2021. Complement factor C1q mediates sleep spindle loss and epileptic spikes after mild brain injury. Science 373:eabj2685.

20. Yu, Z., and B. Morrison, 3rd, 2010. Experimental mild traumatic brain injury induces functional alteration of the developing hippocampus. J. Neurophysiol. 103:499–510.

21. Cramer, S. W., S. P. Haley, L. S. Popa, R. E. Carter, E. Scott, E. B. Flaherty, J. Dominguez, J. D. Aronson, L. Sabal, D. Surinach, C. C. Chen, S. B. Kodandaramaiah, and T. J. Ebner, 2022. Wide-field calcium imaging reveals widespread changes in cortical functional connectivity following mild traumatic brain injury in the mouse. Neurobiol. Dis. 176:105943.

22. Dwyer, M. K. R., I. Polsfuss, K. Herbert, N. Varghese, and B. Morrison, 3rd, 2025. Long-term potentiation and long-term depression are both impaired afterin vitrostretch injury measured with stretchable microelectrode arrays. Biomed. Phys. Eng. Express 11:045029.

23. Zimmerman, K. A., J. Cournoyer, H. Lai, S. B. Snider, D. Fischer, S. Kemp, C. Karton, T. B. Hoshizaki, M. Ghajari, and D. J. Sharp, 2023. The biomechanical signature of loss of consciousness: computational modelling of elite athlete head injuries. Brain 146:3063–3078.

24. Farid, M. H., M. Ramzanpour, M. Ziejewski, and G. Karami, 2018. Estimating the Brain Strain Rates During Traumatic Brain Injury. Biomed. Sci. Instrum. 54 (1):361–368.

25. Carlsen, R. W., A. L. Fawzi, Y. Wan, H. Kesari, and C. Franck, 2021. A quantitative relationship between rotational head kinematics and brain tissue strain from a 2-D parametric finite element analysis. Brain Multiphysics 2:100024.

26. Bar-Kochba, E., M. T. Scimone, J. B. Estrada, and C. Franck, 2016. Strain and rate-dependent neuronal injury in a 3D in vitro compression model of traumatic brain injury. Sci. Rep. 6:30550.

27. Braun, N. J., K. R. Yao, P. W. Alford, and D. Liao, 2020. Mechanical injuries of neurons induce tau mislocalization to dendritic spines and tau-dependent synaptic dysfunction. Proc. Natl. Acad. Sci. U. S. A. 117:29069–29079.

28. Adams, A. A., Y. Li, H. A. Kim, and B. J. Pfister, 2023. Dorsal root ganglion neurons recapitulate the traumatic axonal injury of CNS neurons in response to a rapid stretch in vitro. Front. Cell. Neurosci. 17:1111403.

29. Geddes, D. M., R. S. Cargill, 2nd, and M. C. LaPlaca, 2003. Mechanical stretch to neurons results in a strain rate and magnitude-dependent increase in plasma membrane permeability. J. Neurotrauma 20:1039–1049.

30. Keating, C. E., and D. K. Cullen, 2021. Mechanosensation in traumatic brain injury. Neurobiol. Dis. 148:105210.

31. Sullivan, D., B. J. Vaglio, M. M. Cararo-Lopes, R. D. P. Wong, O. Graudejus, and B. L. Firestein, 2024. Stretch-Induced Injury Affects Cortical Neuronal Networks in a Time- and Severity-Dependent Manner. Ann. Biomed. Eng. 52:1021–1038.

32. LaPlaca, M. C., D. K. Cullen, J. J. McLoughlin, and R. S. Cargill, 2nd, 2005. High rate shear strain of three-dimensional neural cell cultures: a new in vitro traumatic brain injury model. J. Biomech. 38:1093–1105.

33. Geddes-Klein, D. M., K. B. Schiffman, and D. F. Meaney, 2006. Mechanisms and consequences of neuronal stretch injury in vitro differ with the model of trauma. J. Neurotrauma 23:193–204.

34. Monnerie, H., M. D. Tang-Schomer, A. Iwata, D. H. Smith, H. A. Kim, and P. D. Le Roux, 2010. Dendritic alterations after dynamic axonal stretch injury in vitro. Exp. Neurol. 224:415–423.

35. LaPlaca, M. C., and L. E. Thibault, 1998. Dynamic mechanical deformation of neurons triggers an acute calcium response and cell injury involving the N-methyl-D-aspartate glutamate receptor. J. Neurosci. Res. 52:220–229.

36. Belin, S., B. A. Maki, J. Catlin, B. A. Rein, and G. K. Popescu, 2022. Membrane Stretch Gates NMDA Receptors. J. Neurosci. 42:5672–5680.

37. Gu, Y., P. Jukkola, Q. Wang, T. Esparza, Y. Zhao, D. Brody, and C. Gu, 2017. Polarity of varicosity initiation in central neuron mechanosensation. J. Cell Biol. 216:2179–2199.

38. Chi, S., Y. Cui, H. Wang, J. Jiang, T. Zhang, S. Sun, Z. Zhou, Y. Zhong, and B. Xiao, 2022. Astrocytic Piezo1-mediated mechanotransduction determines adult neurogenesis and cognitive functions. Neuron 110:2984–2999.e8.

39. Kanda, H., K. Noguchi, and Y. Dai, 2023. Axonal membrane stretch suppresses neuronal excitability by activating mechanosensitive K2P channels at the node of Ranvier. Mol. Brain 16:8.

40. Magou, G. C., B. J. Pfister, and J. R. Berlin, 2015. Effect of acute stretch injury on action potential and network activity of rat neocortical neurons in culture. Brain Res. 1624:525–535.

41. Hoffe, B., and M. R. Holahan, 2022. Hyperacute Excitotoxic Mechanisms and Synaptic Dysfunction Involved in Traumatic Brain Injury. Front. Mol. Neurosci. 15:831825.

42. Beamer, M., S. R. Tummala, D. Gullotti, C. Kopil, S. Gorka, Ted Abel, C. R. D. Bass, B. Morrison, 3rd, A. S. Cohen, and D. F. Meaney, 2016. Primary blast injury causes cognitive impairments and hippocampal circuit alterations. Exp. Neurol. 283:16–28.

43. Bain, A. C., and D. F. Meaney, 2000. Tissue-level thresholds for axonal damage in an experimental model of central nervous system white matter injury. J. Biomech. Eng. 122:615–622.

44. Wright, R. M., and K. T. Ramesh, 2012. An axonal strain injury criterion for traumatic brain injury. Biomech. Model. Mechanobiol. 11:245–260.

45. Hansen, K. R., G. J. DeWalt, A. I. Mohammed, H.-A. Tseng, M. E. Abdulkerim, S. Bensussen, V. Saligrama, B. Nazer, W. D. Eldred, and X. Han, 2018. Mild Blast Injury Produces Acute Changes in Basal Intracellular Calcium Levels and Activity Patterns in Mouse Hippocampal Neurons. J. Neurotrauma 35:1523–1536.

46. Summey, L., J. Zhang, A. K. Landauer, J. Sergay, J. Yang, A. Daul, J. Tao, J. Park, A. McGhee, and C. Franck, 2023. Open Source, In-Situ, Intermediate Strain-Rate Tensile Impact Device for Soft Materials and Cell Culture Systems. Exp. Mech. 63:1445–1460.

47. Scimone, M. T., H. C. Cramer, III, E. Bar-Kochba, R. Amezcua, J. B. Estrada, and C. Franck, 2018. Modular approach for resolving and mapping complex neural and other cellular structures and their associated deformation fields in three dimensions. Nat. Protoc. 13:3042–3064.

48. Herculano-Houzel, S., and R. Lent, 2005. Isotropic fractionator: a simple, rapid method for the quantification of total cell and neuron numbers in the brain. J. Neurosci. 25:2518–2521.

49. Song, K., N.-K. Cho, K. Park, and C.-S. Kim, 2022. Investigating mechanical behaviours of PDMS films under cyclic loading. Polymers (Basel) 14:2373.

50. Jang, M. J., and Y. Nam, 2015. NeuroCa: integrated framework for systematic analysis of spatiotemporal neuronal activity patterns from large-scale optical recording data. Neurophotonics 2:035003.

51. Sergay, J. A., A. Hai, and C. Franck, 2025. NeuroSpikeX: Comprehensive detection and characterization of neuronal calcium dynamics. SoftwareX 32:102435.

52. Weber, J. T., 2004. Calcium homeostasis following traumatic neuronal injury. Curr. Neurovasc. Res. 1:151–171.

53. Weber, J. T., 2012. Altered calcium signaling following traumatic brain injury. Front. Pharmacol. 3:60.

54. Lusardi, T. A., J. Rangan, D. Sun, D. H. Smith, and D. F. Meaney, 2004. A device to study the initiation and propagation of calcium transients in cultured neurons after mechanical stretch. Ann. Biomed. Eng. 32:1546–1558.

55. Staal, J. A., T. C. Dickson, R. Gasperini, Y. Liu, L. Foa, and J. C. Vickers, 2010. Initial calcium release from intracellular stores followed by calcium dysregulation is linked to secondary axotomy following transient axonal stretch injury. J. Neurochem. 112:1147–1155.

56. Wolf, J. A., P. K. Stys, T. Lusardi, D. Meaney, and D. H. Smith, 2001. Traumatic axonal injury induces calcium influx modulated by tetrodotoxin-sensitive sodium channels. J. Neurosci. 21:1923–1930.

57. Eisner, D., E. Neher, H. Taschenberger, and G. Smith, 2023. Physiology of intracellular calcium buffering. Physiol. Rev. 103:2767–2845.

58. Zhang, Y., M. Rózsa, Y. Liang, D. Bushey, Z. Wei, J. Zheng, D. Reep, G. J. Broussard, A. Tsang, G. Tsegaye, S. Narayan, C. J. Obara, J.-X. Lim, R. Patel, R. Zhang, M. B. Ahrens, G. C. Turner, S. S.-H. Wang, W. L. Korff, E. R. Schreiter, K. Svoboda, J. P. Hasseman, I. Kolb, and L. L. Looger, 2023. Fast and sensitive GCaMP calcium indicators for imaging neural populations. Nature 615:884–891.

59. Giza, C. C., and D. A. Hovda, 2014. The new neurometabolic cascade of concussion. Neurosurgery 75 Suppl 4:S24–33.

60. Deshpande, L. S., D. A. Sun, S. Sombati, A. Baranova, M. S. Wilson, E. Attkisson, R. J. Hamm, and R. J. DeLorenzo, 2008. Alterations in neuronal calcium levels are associated with cognitive deficits after traumatic brain injury. Neurosci. Lett. 441:115–119.

61. Salsabilian, S., Y. Bibineyshvili, D. J. Margolis, and L. Najafizadeh, 2022. Identifying mild traumatic brain injury using measures of frequency-specified networks. J. Neural Eng. 19:056033.

62. Kang, G., E. K. Moo, R. Banton, O. E. Petel, and A. R. Harris, 2025. Cellular mechanisms of traumatic brain injury. NPJ Biol. Phys. Mech. 2:16.

63. Suresh, J., M. Radojicic, L. L. Pesce, A. Bhansali, J. Wang, A. K. Tryba, J. D. Marks, and W. van Drongelen, 2016. Network burst activity in hippocampal neuronal cultures: the role of synaptic and intrinsic currents. J. Neurophysiol. 115:3073–3089.

64. Mayer, A. R., M. V. Mannell, J. Ling, C. Gasparovic, and R. A. Yeo, 2011. Functional connectivity in mild traumatic brain injury. Hum. Brain Mapp. 32:1825–1835.

65. Wang, C., M. E. Costanzo, P. E. Rapp, D. Darmon, D. E. Nathan, K. Bashirelahi, D. L. Pham, M. J. Roy, and D. O. Keyser, 2017. Disrupted gamma synchrony after mild traumatic brain injury and its correlation with white matter abnormality. Front. Neurol. 8:571.

66. Shibasaki, K., K. Ikenaka, F. Tamalu, M. Tominaga, and Y. Ishizaki, 2014. A novel subtype of astrocytes expressing TRPV4 (transient receptor potential vanilloid 4) regulates neuronal excitability via release of gliotransmitters. J. Biol. Chem. 289:14470–14480.

67. Pacheco, J. M., A. Hines-Lanham, C. Stratton, C. J. Mehos, K. E. McCurdy, N. J. Pinkowski, H. Zhang, C. W. Shuttleworth, and R. A. Morton, 2019. Spreading Depolarizations Occur in Mild Traumatic Brain Injuries and Are Associated with Postinjury Behavior. eNeuro 6.

68. Bean, B. P., 2007. The action potential in mammalian central neurons. Nat. Rev. Neurosci. 8:451–465.

69. Parker, E., R. Aboghazleh, G. Mumby, R. Veksler, J. Ofer, J. Newton, R. Smith, L. Kamintsky, C. M. A. Jones, E. O’Keeffe, E. Kelly, K. Doelle, I. Roach, L. T. Yang, P. Moradi, J. M. Lin, A. J. Gleason, C. Atkinson, C. Bowen, K. D. Brewer, C. P. Doherty, M. Campbell, D. B. Clarke, G. van Hameren, D. Kaufer, and A. Friedman, 2022. Concussion susceptibility is mediated by spreading depolarization-induced neurovascular dysfunction. Brain 145:2049–2063.

70. Murphy, T. H., L. A. Blatter, W. G. Wier, and J. M. Baraban, 1992. Spontaneous synchronous synaptic calcium transients in cultured cortical neurons. J. Neurosci. 12:4834–4845.

71. Ali, F., and A. C. Kwan, 2020. Interpreting in vivo calcium signals from neuronal cell bodies, axons, and dendrites: a review. Neurophotonics 7:011402.

72. Ramakrishna, S., B. K. Radhakrishna, A. P. Kaladiyil, N. M. Shah, N. Basavaraju, K. K. Freude, R. P. Kommaddi, and R. S. Muddashetty, 2024. Distinct calcium sources regulate temporal profiles of NMDAR and mGluR-mediated protein synthesis. Life Sci. Alliance 7:e202402594.

73. Basarsky, T. A., S. N. Duffy, R. D. Andrew, and B. A. MacVicar, 1998. Imaging spreading depression and associated intracellular calcium waves in brain slices. J. Neurosci. 18:7189–7199.

74. Dietz, R. M., J. H. Weiss, and C. W. Shuttleworth, 2009. Contributions of Ca2+ and Zn2+ to spreading depression-like events and neuronal injury. J. Neurochem. 109 Suppl 1:145–152.

75. Brini, M., and E. Carafoli, 2011. The plasma membrane Ca²+ ATPase and the plasma membrane sodium calcium exchanger cooperate in the regulation of cell calcium. Cold Spring Harb. Perspect. Biol. 3:a004168–a004168.

76. O’Day, D. H., 2024. Calcium and non-penetrating traumatic brain injury: A proposal for the implementation of an early therapeutic treatment for initial head insults. Biomolecules 14:853.

77. Ahmadzadeh, H., D. H. Smith, and V. B. Shenoy, 2014. Viscoelasticity of tau proteins leads to strain rate-dependent breaking of microtubules during axonal stretch injury: predictions from a mathematical model. Biophys. J. 106:1123–1133.

78. Tang-Schomer, M. D., A. R. Patel, P. W. Baas, and D. H. Smith, 2010. Mechanical breaking of microtubules in axons during dynamic stretch injury underlies delayed elasticity, microtubule disassembly, and axon degeneration. FASEB J. 24:1401–1410.

79. Ahmed, S. M., B. A. Rzigalinski, K. A. Willoughby, H. A. Sitterding, and E. F. Ellis, 2000. Stretchinduced injury alters mitochondrial membrane potential and cellular ATP in cultured astrocytes and neurons. J. Neurochem. 74:1951–1960.

80. Arundine, M., M. Aarts, A. Lau, and M. Tymianski, 2004. Vulnerability of central neurons to secondary insults after in vitro mechanical stretch. J. Neurosci. 24:8106–8123.

81. Dollé, J.-P., A. Jaye, S. A. Anderson, H. Ahmadzadeh, V. B. Shenoy, and D. H. Smith, 2018. Newfound sex differences in axonal structure underlie differential outcomes from in vitro traumatic axonal injury. Exp. Neurol. 300:121–134.

82. Paredes, R. M., J. C. Etzler, L. T. Watts, W. Zheng, and J. D. Lechleiter, 2008. Chemical calcium indicators. Methods 46:143–151.

83. Mira, R. G., R. A. Quintanilla, and W. Cerpa, 2023. Mild traumatic brain injury induces mitochondrial calcium overload and triggers the upregulation of NCLX in the hippocampus. Antioxidants (Basel) 12:403.

84. Paschen, W., and J. Doutheil, 1999. Disturbances of the functioning of endoplasmic reticulum: a key mechanism underlying neuronal cell injury? J. Cereb. Blood Flow Metab. 19:1–18.

85. Weber, J. T., B. A. Rzigalinski, and E. F. Ellis, 2001. Traumatic injury of cortical neurons causes changes in intracellular calcium stores and capacitative calcium influx. J. Biol. Chem. 276:1800–1807.

86. Barykina, N. V., O. M. Subach, K. D. Piatkevich, E. E. Jung, A. Y. Malyshev, I. V. Smirnov, A. O. Bogorodskiy, V. I. Borshchevskiy, A. M. Varizhuk, G. E. Pozmogova, E. S. Boyden, K. V. Anokhin, G. N. Enikolopov, and F. V. Subach, 2017. Green fluorescent genetically encoded calcium indicator based on calmodulin/M13-peptide from fungi. PLoS One 12:e0183757.

87. Chen, T.-W., T. J. Wardill, Y. Sun, S. R. Pulver, S. L. Renninger, A. Baohan, E. R. Schreiter, R. A. Kerr, M. B. Orger, V. Jayaraman, L. L. Looger, K. Svoboda, and D. S. Kim, 2013. Ultrasensitive fluorescent proteins for imaging neuronal activity. Nature 499:295–300.

88. Nakai, J., M. Ohkura, and K. Imoto, 2001. A high signal-to-noise Ca(2+) probe composed of a single green fluorescent protein. Nat. Biotechnol. 19:137–141.

89. Zhong, B. L., V. T. Vachharajani, and A. R. Dunn, 2022. Facile detection of mechanical forces across proteins in cells with STReTCh. Cell Rep. Methods 2:100278.

90. Sergay, J. A., N. Schick, E. Blick, M. Araiza Canizales, J. Zhang, C. Franck, and A. Hai, 2025. Data from: Mechanical strain and strain rate define thresholds of network-level calcium disruption after traumatic brain injury. Dryad Digital Repository, dataset. 10.5061/dryad.1jwstqk9h.

