## Supplemental Material for "Mechanical strain and strain rate define thresholds of single-cell and ensemble calcium disruption in traumatic brain injury"

Contents: Supporting Methods; Supporting Text; Figures S1–S6; Table S1.

### Supporting Methods

#### GCaMP6s indicator saturation analysis

Because samples stretched at 0.5 strain at 50 $s^{-1}$ exhibited a pronounced increase in basal GCaMP fluorescence levels (main text, Fig. 3), we sought to determine whether the calcium indicator approached its saturation limit, which could confound measurements of calcium transients. First, all time-lapse recordings for this condition were reviewed in Nikon Elements to confirm that raw fluorescence intensities were not clipped at the detector level. After confirming the absence of microscope saturation, an empirical, cell-level measure of indicator saturation—termed *headroom*—was computed in MATLAB for all sham and 0.5 strain samples at 50 $s^{-1}$.

$$\text{Headroom} = 1 - \frac{F_{0,i}}{F_{sat}} \tag{S1}$$

Here, $F_{0,i}$ denotes the basal fluorescence of cell $i$, calculated as the 10th percentile of its raw fluorescence trace, and $F_{sat}$ denotes the 95th percentile of all raw fluorescence values within the same recording, serving as an empirical estimate of the upper fluorescence ceiling of the indicator.

To classify cells operating near this ceiling, a low-headroom threshold was defined as the 5th percentile of headroom values pooled across all sham cells at 0+. Cells previously identified as exhibiting elevated basal fluorescence, their spike rates at 0- and 0+, and their headroom classifications were then compared. Culture-level statistical analyses (RStudio) were run by fitting Gaussian linear models with condition (sham vs. stretch) as a fixed effect and treating each culture as an independent replicate. Statistical significance was assessed using two-sided t-tests on model coefficients.

#### Basal fluorescence shift and event rate association

To determine whether hyperacute calcium basal elevations directly corresponded to decreased neuronal activity, we tested whether per-cell increases in basal fluorescence predicted decreases in calcium event rate immediately after stretch for the 0.5 strain at 50 $s^{-1}$ condition. Primary analyses were restricted to neurons that remained active at both 0- and 0+ time points.

For each culture, cells were classified as exhibiting elevated basal fluorescence when the $log_2$ fold-change exceeded the predefined threshold (main text, Materials and Methods: Neuronal calcium transient assessment). Meaningful calcium event-rate reduction was set to $\leq$-60%. It was defined similarly by identifying the middle 95th percentile of $\Delta$ER across all sham samples, with a decrease $\geq$60%.

Associations were tested using a mixed-effects logistic regression to account for clustering within cultures. A stratified Cochran–Mantel–Haenszel (CMH) test was also performed as a non-parametric confirmation. As a secondary analysis, basal elevation was calculated as a predictor of the immediate loss of calcium activity (from active at 0- to inactive at 0+) across the full cohort.

#### Spatial network analysis

To characterize injury-induced changes in functional connectivity, pairwise Pearson correlations between the smoothed calcium fluorescence signals of all simultaneously recorded active cells were computed per sample and time point. A correlation threshold was calibrated data-adaptively by sweeping candidate values from 0 to 0.99 and selecting the threshold at which the mean fraction of connected cell pairs across all pre-injury (0-) recordings equaled 0.50, providing a balanced starting point from which injury-induced changes could be detected in either direction. The fraction of connected cell pairs was defined as the number of cell pairs with pairwise correlation above this threshold divided by the total number of possible pairs. Mean degree and mean local clustering coefficient were computed from the resulting binary adjacency matrix. Spatial autocorrelation of calcium activity was quantified using Moran's $I$ with an inverse-distance spatial weight matrix; statistical significance was assessed via a 999-permutation test. Cell soma positions were extracted by NeuroCa and converted to micrometers using a calibrated pixel size of 1.28 $\mu$m/pixel (Nikon Ti-E, 10×/0.45 NA, Kinetix camera, 2×2 binning).

Network connectivity metrics were analyzed using linear mixed-effects models (LME; `lme4` and `lmerTest` packages, R) fitted by restricted maximum likelihood, with SampleID as a random intercept to account for repeated measures across time points within samples. Strain condition and time point were included as fixed effects with their interaction. Control cultures, which serve only for the device- and imaging-control comparison shown in Fig. S2, were excluded from these models, so that the sham condition is the reference throughout. Post-hoc pairwise contrasts within conditions were Tukey-adjusted, and comparisons of each stretch condition against sham were Dunnett-adjusted, using the `emmeans` package. Degrees of freedom were approximated by Satterthwaite's method. Statistical significance was defined as $p < 0.05$.

#### Leave-one-out sensitivity analysis

To assess the robustness of each significant finding to individual sample contributions, a leave-one-out (LOO) sensitivity procedure was applied to each primary outcome reported in the main text. For each finding, the model used in the primary analysis was refit $N$ times, where $N$ is the number of independent cultures contributing to that finding, omitting one culture per refit. The same family, link, fixed effects, and random-intercept structure as the primary model were used for every refit. Convergence and Hessian positive-definiteness were verified for each refit, and refits that failed either check were flagged. For each finding, the number of refits retaining $p < 0.05$ and the maximum $p$ value observed across refits were recorded, and the result was classified as *Robust* (no refit lost significance) or *Review* (at least one refit lost significance). Results across all key findings are summarized in Table S1.

### Supporting Text

#### Device contact and imaging do not change calcium dynamics

To isolate the effects of the injury device and imaging protocol, we recorded GCaMP6s activity and Hoechst/Ethidium signals in "control" cultures that were never placed in the tension device and compared them with "sham" cultures that were mounted but not stretched. Two-minute recordings were segmented to mimic 0- and 0+ time windows.

Within-group changes from 0- to 0+ were not significant for either calcium event rate or burst rate in Control or Sham (Fig. S2A–B; Tweedie GLMM throughout). Event rate: Control $\Delta$ER $= 44.76\%$, 95% CI $[-12.94, 140.69]\%$, $N = 5$, $p = 0.155$; Sham $\Delta$ER $= 2.31\%$, 95% CI $[-27.48, 44.35]\%$, $N = 10$, $p = 0.896$. Burst rate: Control $\Delta$Burst Rate $= -18.48\%$, 95% CI $[-38.79, 8.56]\%$, $p =$

0.162; Sham $\Delta$Burst Rate $= 1.26\%$, 95% CI $[-14.84, 20.40]\%$, $p = 0.888$. Baseline (0-) levels were also comparable: $\Delta ER_{0-}$(Control–Sham) $= -22.59\%$, 95% CI $[-61.72, 56.55]\%$, $p = 0.476$; $\Delta$Burst Rate$_{0-}$(Control–Sham) $= -52.79\%$, 95% CI $[-85.19, 50.49]\%$, $p = 0.204$.

In these controls, we acquired Hoechst and Ethidium once at 24h, so Eth$^+$ reflects membrane permeation at 24 h but cannot be used to compute matched per-nucleus death, as in the main text (which requires aligned Hoechst at 0- and 24h). Using this partial metric (beta-binomial model, Tukey-adjusted across all groups), the fraction of Eth$^+$ nuclei was significantly lower in Control than Sham (difference $= -7.28$ percentage points, 95% CI $[-11.90, -2.65]$, $p = 5.1 \times 10^{-5}$), and lower than $\varepsilon = 0.3$ @ $1\,s^{-1}$ (difference $= -7.34$ percentage points, 95% CI $[-14.10, -0.58]$, $p = 0.022$)—noting higher variability in that condition. $\varepsilon = 0.3$ @ $50\,s^{-1}$ was modestly lower than Sham (difference $= -5.41$ percentage points, 95% CI $[-10.73, -0.10]$, $p = 0.042$). These Eth$^+$ differences indicate the mount/imaging protocol introduces a small amount of membrane permeation relative to untouched controls. However, consistent with the main-text death metric that tracks nuclei between 0- and 24h, stretch conditions did not show increased cell death compared with Sham. Thus, the network-level changes reported here are not explained by device contact or imaging and are not driven by widespread cell loss.

### Increased basal calcium fluorescence levels after severe stretch do not confound network hypoactivity

Across cultures, stretch significantly increased the fraction of low-headroom cells relative to sham ($\beta = 0.084 \pm 0.034$; $N = 5 - 10$ cultures per condition; $p = 0.026$), indicating that a greater proportion of cells operated near the upper dynamic range of GCaMP following injury (Fig. S3A). Critically, even after excluding all low-headroom cells, stretch cultures exhibited a significantly larger reduction in calcium event rate (median $\Delta$ ER, 0+ vs 0-) compared with shams ($\beta = -0.060 \pm 0.016$ Hz; $N = 5$–$10$; $p = 0.0027$), supporting a genuine biological reduction of network activity rather than a measurement artifact (Fig. S3B).

The difference in headroom between cells identified as having elevated basal fluorescence and those with no basal change did not differ significantly by condition ($p = 0.083$). As illustrated by a representative 0.5 strain at 50 $s^{-1}$ sample (Fig. S3C), these two cell populations were interspersed across the full range of fluorescence headroom. Within this network, event rate at 0+ showed no meaningful correlation with headroom (Spearman $\rho = 0.078$, $p = 0.149$, slope $= 0.0025$), and the change in event rate from 0- to 0+ exhibited only a weak association with negligible effect size ($\rho = 0.111$, $p = 0.0395$, slope $= 0.006$). Together, these results indicate that reduced calcium transient activity following stretch is not driven by proximity to the GCaMP saturation ceiling.

### Cell-level cytosolic calcium changes do not predict decreased calcium transients

We asked whether neurons exhibiting elevated basal fluorescence immediately after stretch, representative of increased cytosolic calcium, were the same neurons that reduced calcium transient activity. Because neurons that became inactive at 0+ undergo a fixed -100% change in event rate, analyses were restricted to cells that remained active throughout the hyperacute transition to isolate modulation of activity from loss of activity. Across individual neurons, basal fluorescent elevation showed no relationship with calcium event rate reduction (Fig. S6A). Cells exhibited a wide range of changes in event rates regardless of baseline calcium event state, and no consistent trend was observed across cultures.

This relationship was tested statistically using a mixed-effects logistic regression model that accounted for culture-to-culture variability. Elevated basal fluorescence did not predict calcium event reduction among active neurons (Fig. S6B; OR = 0.99, p = 0.96; CMH p = 0.95). Consistent with this result, the probability of calcium event decrease was similar for neurons with and without elevated basal fluorescence within each culture (Fig. S6C). As a secondary analysis, basal elevation also did not reliably predict immediate loss of activity (OR = 1.17, p = 0.14).

These findings demonstrate that hyperacute elevation of the cytosolic calcium and reduction of calcium transients occur independently at the single-cell level, indicating parallel injury responses rather than a shared cellular mechanism at higher levels of applied stretch.

### Network burst activity and neuronal participation across strains and time points

To provide additional context for network-level dynamics following stretch, we examined burst activity and neuronal participation across strains and time points (Fig. S4). At the slower strain rate (1 $s^{-1}$), network burst rates remained broadly similar across strains and time points, with only a small number of comparisons reaching statistical significance (Fig. S4A). At the faster strain rate (50 $s^{-1}$), burst activity showed greater variability across strains, including a significant reduction immediately after stretch at $\varepsilon$ = 0.3 and loss of bursting in every culture at $\varepsilon$ = 0.5 (Fig. S4B). At 24h, network bursts were absent in 5 of 10 sham cultures and in 4 of 5 cultures stretched to $\varepsilon$ = 0.5 at 50 $s^{-1}$.

We also quantified the number of active neurons across strains at the slower strain rate (Fig. S4C). Across these conditions, the number of active cells remained relatively stable over time, except at $\varepsilon$ = 0.3, where it declined by 24h. Together, these results provide additional context for the main-text analyses by showing how burst activity and neuronal participation evolve across loading conditions and time points.

### Functional network connectivity is selectively disrupted by high-rate, high-magnitude stretch

To complement cell-level calcium metrics, we characterized the spatial organization of functional connectivity across the recorded network. The interaction between strain condition and time point was not significant for the fraction of connected cell pairs ($F(12, 64.3) = 1.61$; $p = 0.112$; LME). The comparisons below are therefore planned within-condition contrasts, motivated by the loss of network bursting already established at $\varepsilon = 0.5$ @ $50\,s^{-1}$, rather than post hoc tests conditioned on an omnibus result (Fig. S5A). Among all conditions tested, only $\varepsilon = 0.5$ @ $50\,s^{-1}$ showed a within-group reduction: the fraction of connected cell pairs collapsed acutely after stretch (0- vs. 0+: estimate $= -0.437 \pm 0.130$; $t_{(63.1)} = -3.32$; $N = 5$; $p = 0.004$, Tukey-adjusted LME) and remained suppressed at 24 hours (0- vs. 24h: estimate $= -0.469 \pm 0.130$; $t_{(63.1)} = -3.56$; $N = 5$; $p = 0.002$, Tukey-adjusted LME), with no recovery between the acute and delayed time points (0+ vs. 24h: $p = 0.97$). At 0+, this condition also showed lower connectivity than sham cultures (estimate $= -0.441 \pm 0.145$; $t_{(74.6)} = -3.03$; $N = 5$; $p = 0.018$, Dunnett-adjusted LME); the corresponding contrast at 24 hours did not reach significance ($t_{(76.3)} = -2.60$; $p = 0.056$). No other condition showed a significant change in connectivity relative to baseline or sham at any time point. A beta regression fitted to the same data, which respects the bounded nature of a proportion, reproduced both within-condition results. Moran's $I$ spatial autocorrelation was uniformly small across all conditions and time points and showed no significant variation by strain condition, time point, or their interaction (all $p > 0.82$; LME), indicating that mechanical stretch does not produce spatially patterned suppression of network activity (Fig. S5B).

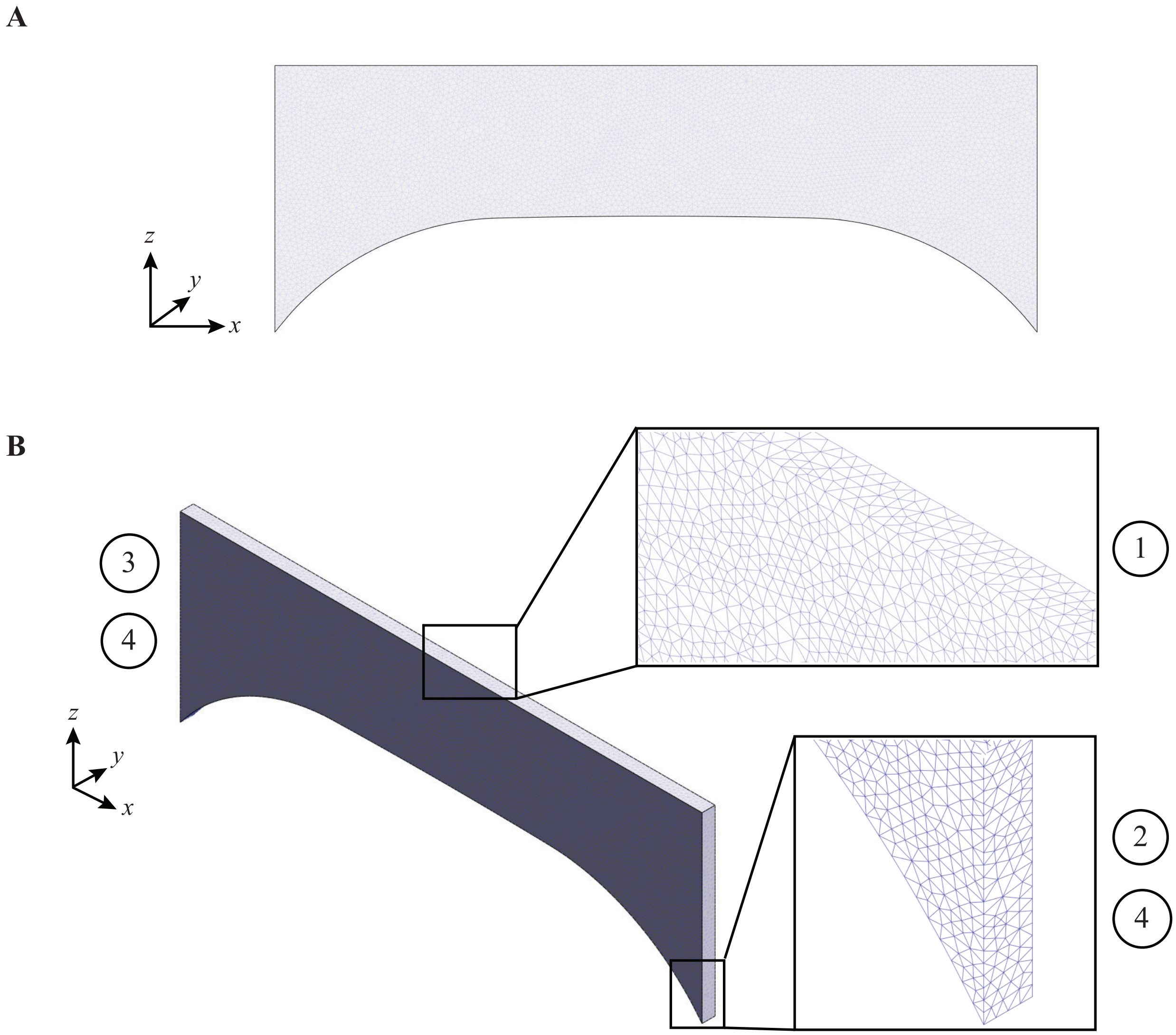


**Figure S1. Dog-bone geometry and boundary conditions for finite element simulations.** (**A**) Front view of the half-symmetry geometry of the dog-bone's gauge region used for finite element simulations. (**B**) Isometric view of the dog-bone geometry with insets highlighting the mesh around the half-symmetry plane and on the fixed ends. The labeled boundary conditions (BC) include: (1) *Symmetry BC* zero displacement in y- and z-direction, (2) *Displacement BC* +2.5 mm displacement in x-direction, (3) *Displacement BC* -2.5 mm displacement in x-direction, and (4) *Fixed-Ends BC* zero displacement in y- and z-direction.

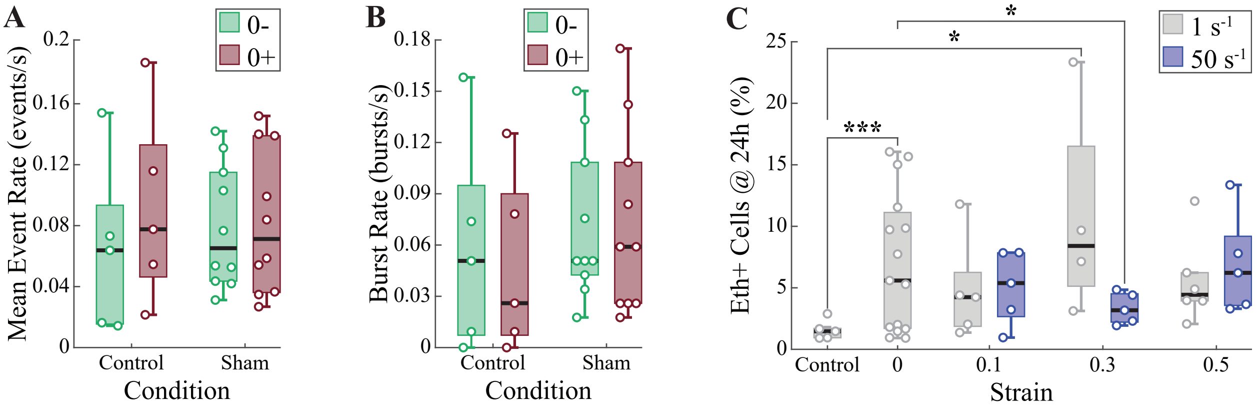


**Figure S2. Controls confirm the device/imaging protocol does not drive calcium changes.** (**A**) Mean calcium event rate in the first two minutes (0-) vs. last two minutes (0+) for control (never mounted) and sham (mounted, no stretch) groups (box plots; $N = 5-10$). Neither group changes significantly. (**B**) Burst rate within groups for the same comparison and samples as in (A); no significant changes. (**C**) Fraction of Eth$^+$ nuclei at 24h in control, sham, and stretch conditions (box plots; $N = 4-10$); this panel reflects membrane permeation only, unlike the main-text death metric, which uses aligned 0- and 24h Hoechst together with Eth$^+$. Statistics: n.s., $*p < 0.05$, $***p < 0.001$ (see main text, Materials and Methods).

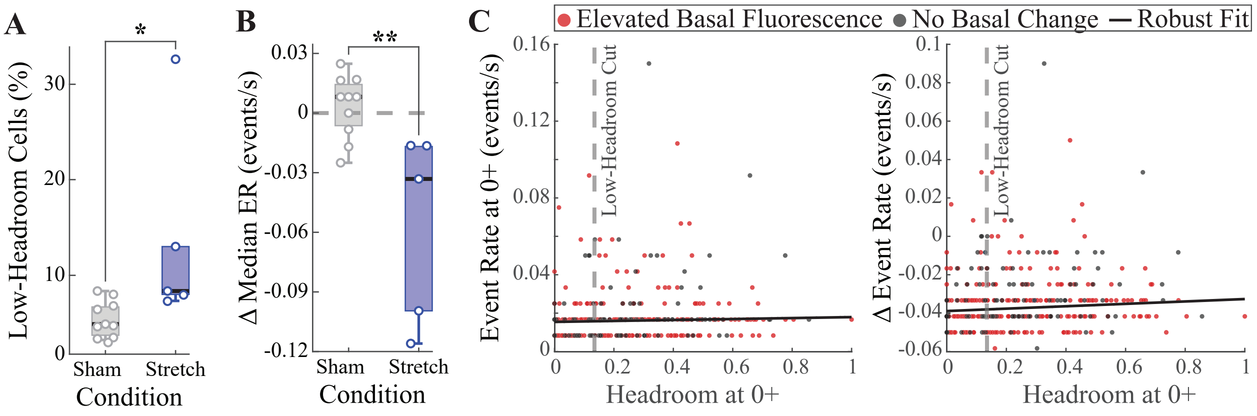


**Figure S3. The observed reduction in calcium transients immediately after stretch is not an artifact of GCaMP over-saturation.** (**A**) Fluorescence headroom was calculated per cell to assess the proximity of basal fluorescence to the usable dynamic range of GCaMP ($Headroom = 1 - F_{0,i}/F_{sat}$; see Supporting Methods). Box plots show the fraction of cells classified as low-headroom in each sham and 0.5 strain at 50 s$^{-1}$ (stretch) sample, based on a sham-derived threshold ($N = 5-10$ cultures per condition). (**B**) After excluding low-headroom cells, stretch cultures still exhibit a greater reduction in the median calcium event rate (0+ vs 0-) than shams ($N = 5-10$ cultures per condition). (**C**) Per-cell analysis from a representative stretch sample relating fluorescence headroom at 0+ to calcium event rate at 0+ (left) and to the change in event rate (0+ vs 0-; right), with cells colored by basal fluorescence shift classification. All statistics: $*p < 0.05$, $**p < 0.01$ (see Supporting Methods).

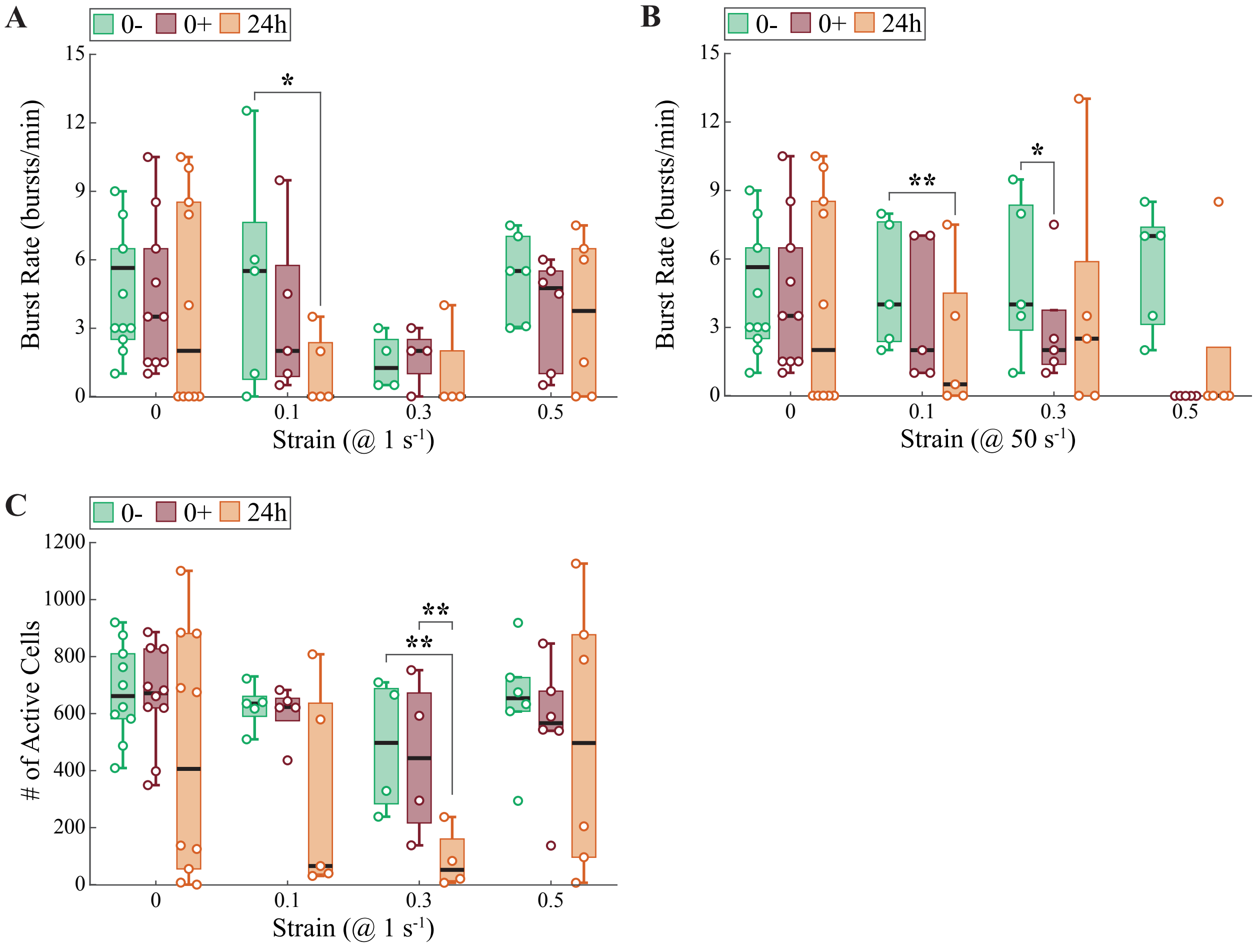


**Figure S4. Network burst activity and neuronal participation across strains and time points.** (**A**) Network burst rate (bursts/min) at baseline (0-), immediately after stretch (0+), and 24 hours later (24h) across strain magnitudes at 1 $s^{-1}$ (box plots; $N = 4 - 10$ cultures per condition). (**B**) Network burst rate across strain magnitudes at 50 $s^{-1}$ for the same time points as in (A) (box plots; $N = 5 - 10$ cultures per condition). (**C**) Number of active cells at 0-, 0+, and 24h across strain magnitudes at 1 $s^{-1}$ (box plots; $N = 4 - 10$ cultures per condition). Individual points represent culture-level measurements. All statistics: $*p < 0.05$, $**p < 0.01$ (Tweedie GLMM; 0+ vs 0- contrasts from the multi-group model and 24 h contrasts from within-condition models; see Methods). In (B), no culture stretched to $\varepsilon = 0.5$ at 50 $s^{-1}$ had network bursts at 0+, so this condition is shown descriptively; see the main text's Fisher's exact test.

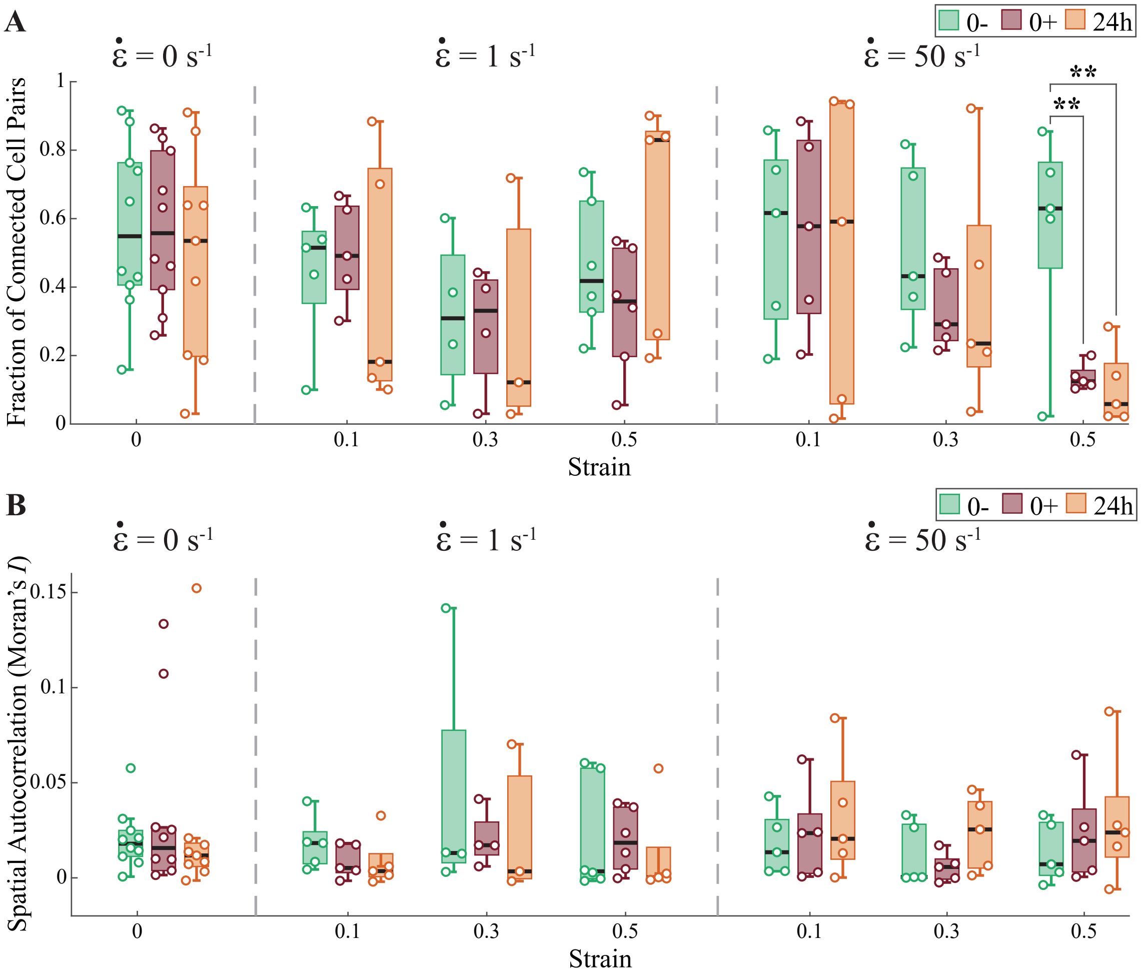


**Figure S5. Spatial network connectivity is selectively disrupted by high-rate, high-magnitude stretch.** (**A**) Fraction of connected cell pairs across strain magnitudes and strain rates at 0-, 0+, and 24h (box plots; $N = 4$–$10$ cultures per condition). A correlation threshold was calibrated so that the mean fraction of connected cell pairs across all pre-injury (0-) recordings equaled 0.50 (see Supporting Methods). (**B**) Spatial autocorrelation (Moran's $I$) of binary cell activity across the same conditions and time points (box plots; $N = 4$–$10$ cultures per condition). Values near zero indicate no spatial clustering of active cells. Statistics: $^{**}p < 0.01$ vs. 0- within group (Tukey-adjusted LME). No significant differences were detected in panel B (all $p > 0.82$, LME).

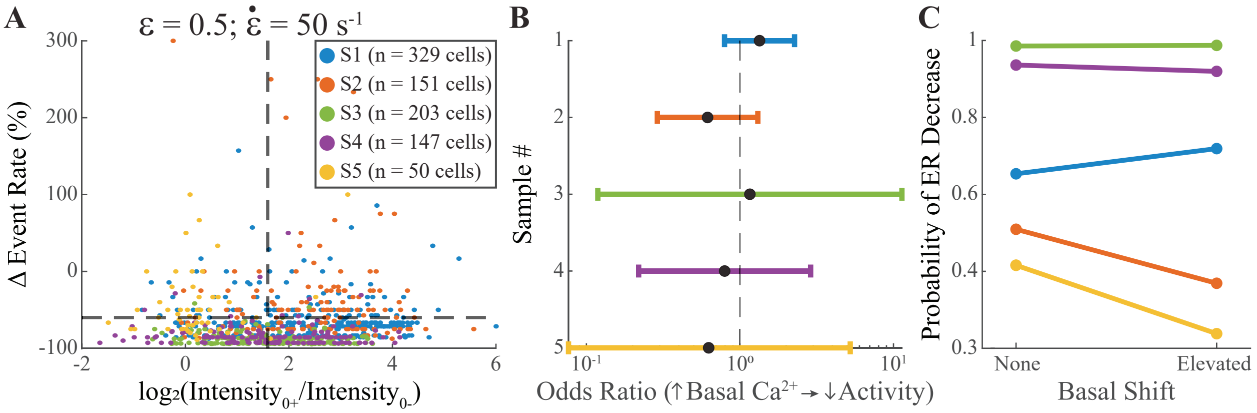


**Figure S6. Hyperacute basal calcium elevation does not predict event-rate suppression.** (**A**) All $\varepsilon$ = 0.5 at 50 $s^{-1}$ samples (0+ vs. 0-): per-cell basal fluorescence change $\left(\log_2(\text{Intensity}_{0+}/\text{Intensity}_{0-})\right)$ versus change in calcium event rate. Points are individual cells; only neurons active at both time points are shown. The vertical dashed line marks the threshold for meaningful basal fluorescence elevation, and the horizontal dashed line marks the threshold for meaningful event-rate reduction. (**B**) Per-culture odds ratios for increased basal fluorescence predicting transient reduction among active neurons. Points represent culture-level estimates with 95% confidence intervals (log scale). The dashed line indicates no association (OR = 1). (**C**) Within-culture probabilities of a meaningful event rate (ER) decrease for cells without versus with elevated basal fluorescence (paired points joined by lines). See Supporting Methods for definitions and analysis details.

**Table S1. Leave-one-out (LOO) sensitivity summary for key findings.** For each significant finding the table reports the full-model effect and $p$-value, the number of LOO refits in which significance was lost out of the total ($n_{\text{lost}}/n$), the maximum $p$-value observed across LOO refits, and the resulting robustness classification. Immediate (0+ vs 0-) and between-condition contrasts were extracted from the multi-group models used for the primary analysis; 24 h contrasts were extracted from the within-condition models fitted separately for each condition (see main text, Materials and Methods). All LOO refits converged with positive-definite Hessians. See Supporting Methods for procedural details.

| Finding | Effect | $p$ | $n_{\text{lost}}/n$ | Max LOO $p$ | Robustness |
|---|---|---|---|---|---|
| Event rate, $\varepsilon$ = 0.3 @ 50 $s^{-1}$ (0+ vs 0-) | −38.05% | $1.2\times10^{-3}$ | 0/40 | 0.021 | Robust |
| Event rate, $\varepsilon$ = 0.5 @ 50 $s^{-1}$ (0+ vs 0-) | −72.07% | $4.0\times10^{-16}$ | 0/40 | $<10^{-10}$ | Robust |
| Rate constant ($k$), $\varepsilon$ = 0.5 @ 50 $s^{-1}$ (0+ vs 0-) | +186.46% | $1.05\times10^{-10}$ | 0/40 | $<10^{-6}$ | Robust |
| Active cells, $\varepsilon$ = 0.5 @ 50 $s^{-1}$ (0+ vs 0-) | −29.43% | 0.005 | 3/40 | 0.069 | Review |
| Elevated basal fluorescence, $\varepsilon$ = 0.5 @ 50 $s^{-1}$ vs Sham (0+) | +37.92 pp | $7.3\times10^{-7}$ | 0/40 | $<10^{-4}$ | Robust |
| Active cells, $\varepsilon$ = 0.1 @ 50 $s^{-1}$ (24 h vs 0-) | −45.13% | 0.020 | 0/5 | 0.037 | Robust |
| Active cells, $\varepsilon$ = 0.3 @ 50 $s^{-1}$ (24 h vs 0-) | −56.23% | 0.006 | 1/5 | 0.066 | Review |
| Active cells, $\varepsilon$ = 0.5 @ 50 $s^{-1}$ (24 h vs 0-) | −81.10% | $1.7\times10^{-6}$ | 0/5 | $<10^{-3}$ | Robust |
| Active cells, $\varepsilon$ = 0.5 @ 50 $s^{-1}$ (24 h vs 0+) | −73.22% | $3.2\times10^{-4}$ | 0/5 | 0.005 | Robust |
| Cell death, $\varepsilon$ = 0.3 (50 $s^{-1}$ vs 1 $s^{-1}$, 24 h) | −12.01 pp | 0.018 | 3/40 | 0.120 | Review |
